# A strategy for genome-wide seamless tagging of human protein-coding genes

**DOI:** 10.1101/2025.03.04.641506

**Authors:** Lidice González, Minoo Karimi, Xavier Castellanos-Girouard, Masih M. Saber, Nicolas Stifani, Louis Gauthier, Adrian W. Serohijos, Stephen W. Michnick

## Abstract

Comprehensive and systematic proteome-wide experiments require financial and technical resources unavailable to most researchers. Here we describe a universal and scalable method based on CRISPR/Cas9-non-homologous end joining (NHEJ) for Pooled Recombinant Integration of Seamless Markers (PRISM) into protein coding genes. We created two genome-wide gRNA libraries for 5’- and 3’-tagging of 18,489 human protein-coding genes. Selection for in-frame integration of the donor cassette can be favored by fusing it to an antibiotic resistance enzyme (ARE) and P2A self-cleaving peptide, resulting in tagged proteins and free ARE to select for clones with inserted donor cassettes. We achieved a library integration rate of ∼20% and tagging of at least 36% of genes expressed in HEK 293T cells including, at least 28% of essential genes, with donor DNA. Our strategy is universal, scalable, specific, and selective, paving the way for genome-scale construction of human cell lines tagged with different types of reporter genes for protein functional characterization.

**Significance Statement:** Here we report a practical method to achieve 5’- or 3’-end integration of reporter protein-coding sequences to express seamless fusions of human protein and reporter proteins that we call **P**ooled **R**ecombinant **I**ntegration of **S**eamless **M**arkers (**PRISM**). PRISM is independent of homology templates, yet it works with high efficiency and precision. Using gRNA-directed CRISPR/Cas9 genome editing and mini-plasmid donor cassettes, we were able to tag at least 36% of the protein- coding sequences of human genes with a green fluorescent protein at an integration efficiency of about 20% in the human cell line HEK 293T. This included tagging approximately 28% of essential genes, an important fact considering their relevance for gene characterization in different cell types.

## Main Text

CRISPR-Cas9 gene editing has revolutionized modern genetics, allowing researchers to precisely knock out genes and regulate their expression at the transcriptional level (1–5). This technology has democratized genetic research, making gene ablation and epistatic analyses widely accessible across laboratories. However, while CRISPR-based gene editing is now routine, large-scale protein tagging remains a challenge. A streamlined approach to tagging proteins would facilitate *in vivo* tracking of protein localization, abundance, and interactions, greatly enhancing functional genomics (6–8). Currently, efforts to create cell libraries with tagged proteins are limited by technical hurdles.

Most gene knock-in existing methods depend on homology-directed repair (HDR), which has low efficiency and requires expensive and time-consuming construction of donor DNA with precise homology arms for each of the ∼20,000 human protein-coding genes (9). As a result, protein-tagging strategies have been applied to only a limited number of genes and cell lines, restricting their broader adoption. Optimization of methods dependent on homology arms include the use of split fluorescent tags (6, 8, 10). For example, the tagging strategy employed for the OpenCell project is based on a split-GFP system, where a small 16 aa C-terminal GFP fragment is inserted endogenously, complemented by an independently expressed N-terminal fragment, allowing systematic mapping of ∼1,300 human protein co-localizations and protein-protein interactions (6, 8). These approaches, although they highly reduce the size of the donor cassette, still rely on the synthesis of specific homology arms, and on HDR low efficiency. Moreover, these strategies are best suited for insertion of small tags, such as half-fragments of reporter proteins, where the other half can be expressed separately. While these techniques provide for excellent seamless tagging of protein-coding genes, expense and low efficiency preclude their use for large-scale tagging applications.

Alternatives to HDR-based methods have improved the scalability, simplicity and reduced cost. These alternatives mainly rely on the integration of the tag cassette through non-homologous end joining repair (NHEJ) at the double strand break, which is much more efficient than HDR, and does not require flanking homology sequences. In this way, only a generic donor cassette is needed containing the desired tag (11–18). For instance, a strategy designed to tag specific gene exons include the use of a generic cassette in a plasmid containing the tag flanked by sgRNA recognition sites targeted by Cas9 (12). This strategy not only guarantees that a small donor cassette is integrated by NHEJ on the targeted exon but also avoids the insertion of non-desired sequences in the genome such as plasmid backbones. This study showed that despite the approach relying on NHEJ, many of the clones obtained contained the tag integrated as designed (12). Nonetheless, since NHEJ repair is prone to errors including frameshift mutations, prior approaches developed circumvented this by targeting intronic sequences (13, 15–17). In this case, homology-independent CRISPR-Cas9 approaches inserted tags as synthetic exons into gene intronic regions (13, 15–17). In addition to the coding sequence for a reporter fluorescent protein and flanking recognition sgRNA sites, the synthetic exons are also flanked by splice donor and acceptor sequences to allow splicing of the tag into one or several introns of a gene of interest (13, 15–17)

Despite these advances, to tag exons of genes themselves more efficiently, strategies are needed to overcome the errors caused by NHEJ. In particular, CRISPaint achieved seamless and in-frame 3’ tag fusions of a donor cassette containing the coding sequence of a tag of interest followed by that for a self-cleaving peptide and an antibiotic resistance enzyme (14). Consequently, clones expressing seamless and in-frame protein products of these fusions can be selected for with an antibiotic, while the self-cleaving peptide separates the tagged protein of interest from the antibiotic resistance enzyme. CRISPaint, although efficient, is designed for tagging one gene at a time. To address this limitation Kim *et al.* developed a variation of this method: the High-throughput Insertion of Tags Across the Genome (HITAG), allowing efficient pooled tagging of proteins at their C-termini including 244 stress granule–associated proteins and 531 transcription factors (11). However, more scalable and generalizable proteome-wide tagging approaches are still needed for various functional genomic applications.

Weighing the features of the methods described above, we found that optimizing another variation of the CRISPaint strategy would be ideal for large-scale tagging of genes. As described by Leonetti *et al.*(6), an effective strategy for large-scale endogenous protein tagging should meet four fundamental criteria: (i) scalability, facilitating the efficient and cost-effective tagging of numerous genes; (ii) specificity, ensuring precise insertion of tags at the target genomic locus without incorporating extraneous sequences such as selection markers or plasmid backbones; (iii) versatility of the tag, allowing for different analyses; and (iv) selective enrichment of successfully modified cells. We add to this list, (v) universality, allowing a single donor template to be applicable for tagging diverse genomic loci in different cell types. Here, we introduce a CRISPR- Cas9-based approach for the seamless integration of DNA cassettes into human protein-coding genes at a near-proteome-wide scale. This method, termed Pooled Recombinant Integration of Seamless Markers (PRISM), is broadly applicable to dividing cells, technically accessible to researchers, and satisfies all the specified criteria described above, making it a powerful tool for functional proteomics.

## Results

### Overall strategy

PRISM relies on six key elements, some shared with CRISPaint and HITAG, but others unique to PRISM: **i**, a CRISPR/Cas9 guide RNA directed nuclease SpCas9-NG, which has an expanded repertoire of protospacer-adjacent motif (PAM) specificities (19–21); **ii**, a lentiviral sgRNA library; **iii**, the highly efficient non-homologous end joining (NHEJ) repair mechanism; **iv**, a universal one- cut minicircle DNA plasmid to deliver donor DNA into the cells; **v**, a unique “Scramble” gRNA (Sc-gRNA) that directs Cas9 for highly efficient cleavage of the donor plasmid within cells (18); **vi**, a promoterless porcine teschovirus-1 (P2A) self-cleaving peptide (22) fused to an antibiotic resistance gene to select for cells which have the antibiotic resistance gene integrated in-frame (Fig. 1).

**Fig. 1:**
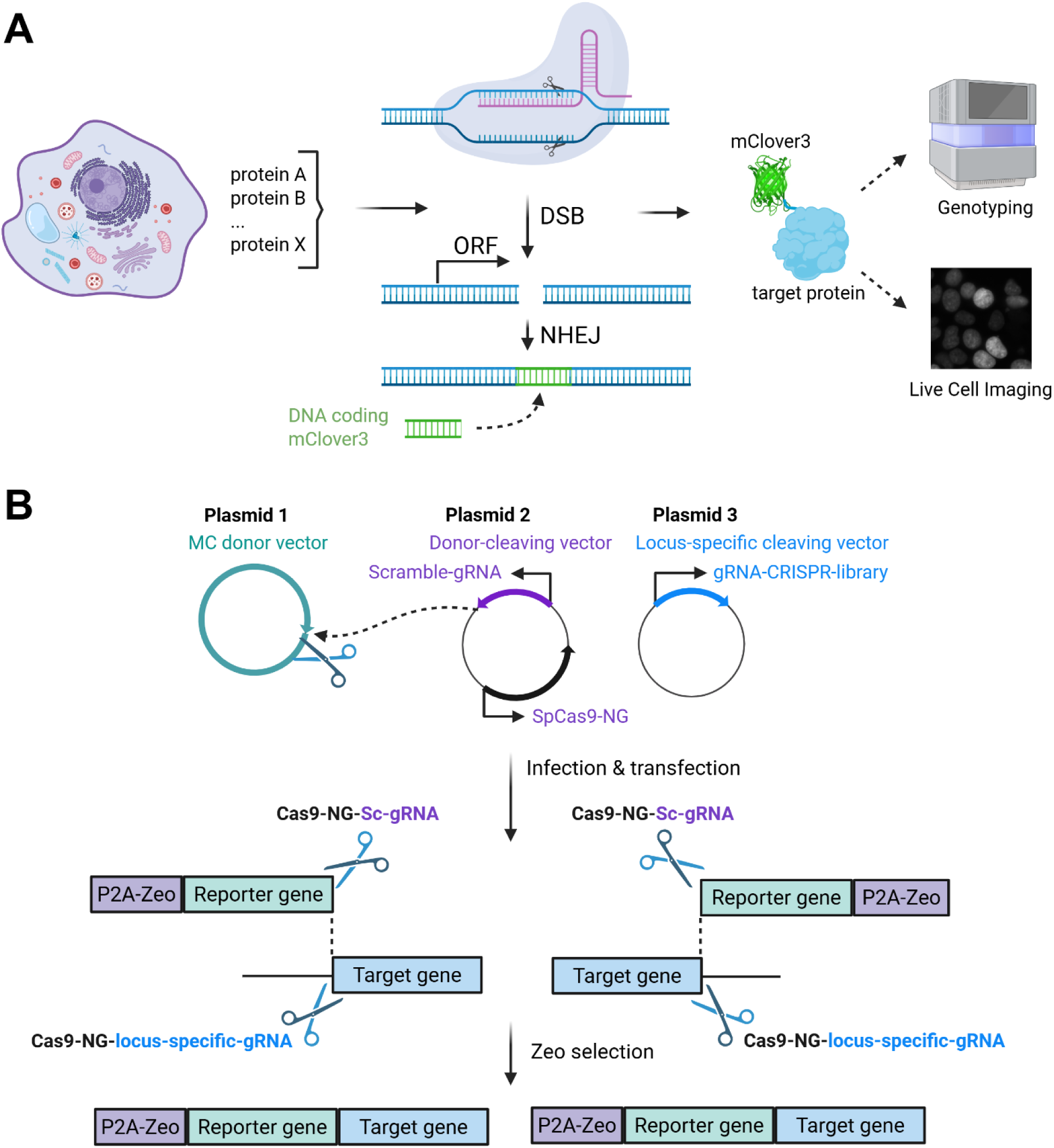
Design of CRISPR/Cas9-non-homologous end joining (NHEJ) based method for Pooled Recombinant Integration of Seamless Markers (PRISM). (A) Overview of the CRISPR-Cas9 editing strategy to seamlessly integrate DNA cassettes as fusions to human protein-coding genes and ways in which the results have been evaluated. (B) Schematic of knock-in vectors and overview of gene editing. The system comprises three plasmids: plasmid 1, a “donor vector” that encodes a promoterless cassette including an antibiotic resistance gene as a selection marker fused to P2A self-cleaving peptide (purple); plasmid 2, a “donor cleaving vector” that carries the Scramble-gRNA (Sc-gRNA) (purple) and Cas9 variant SpCas9-NG (black); and plasmid 3, a lentiviral library of “locus-specific vector” that encodes gRNAs targeting 5’ and 3’ ends of protein-coding sequences of 18,489 human genes that defines the genomic target to be modified by SpCas9-NG (blue). Created in BioRender.com.

Broadly, CRISPR/Cas9-based genome editing begins with the RNA-guided endonuclease Cas9 introducing DNA double-strand breaks at genomic target sites. These double-strand breaks (DSB) then trigger three main categories of DNA repair mechanisms: homology-directed repair (HDR), non-homologous end joining (NHEJ), and microhomology- mediated end joining (MMEJ); each with its own advantages and disadvantages for gene editing (23–27). Although HDR and MMEJ lead to precise repair, their rate is low because they occur only during late S to G2 phases (9, 24, 25, 27). Construction of targeting vectors with the required left and right homology arms (0.5-1 kb) is also cost prohibitive when applied at a large scale. HDR and MMEJ have, however, been successfully applied in several gene knock-in studies, where the low repair rate is offset by increasing the length of homology arms or inhibiting the NHEJ pathway (28–31). Although the NHEJ repair rate is significantly higher than HDR, many events result in unproductive out-of-frame or mutant transcripts and thus, NHEJ is known as an error prone repair system (7, 32). To decrease the probability of errors that could occur by NHEJ, we took advantage of the donor design developed by Schmid-Burgk *et al.* (14), and generated promoter-less integration cassettes consisting of coding sequences for an antibiotic resistance gene (Zeo) fused to a P2A self-cleavage peptide at either its 5’- or 3’-end, and a reporter gene, the green fluorescent protein variant mClover3 (33). P2A self-cleaves at a rate of ∼96% resulting in protein fusions that are dissociated from the antibiotic resistance enzyme (22). Additionally, using antibiotic selection, only cells in which the antibiotic resistance enzyme was integrated in-frame should survive (14). Although the design of the donor cassette highly favors in-frame integration for 3’ tagging, in-frame 5’ integration is not guaranteed by this strategy, so secondary readouts, such as visualization of fused fluorescent protein, provides a secondary selection for in-frame fusions (Fig. 1B).

By using this approach, cells with NHEJ events containing in-frame codon insertions and deletions (indels) could still survive selection, but it is unlikely these would alter the stability or function of proteins more than the fusions themselves. Furthermore, we perform 5’ and 3’ integrations so that either N- or C-terminal fusion proteins are created, providing an extra selection for fusions that do not affect the expression, stability, or function of proteins.

Finally, the use of single-cut minicircle plasmid (MC) to deliver donor DNA is another advantage of this strategy given its improved integration efficiency and low toxicity due to its small size and increased lifetime in cells (18, 34). Furthermore, MCs lack bacterial backbone sequences, eliminating undesired potential integration of backbone plasmid. Overall, PRISM provides a robust means to integrate any large donor DNA cassette into protein-coding genes at near genome-wide scale with product proteins seamlessly fused to a reporter protein of interest.

## Design of seamless DNA cassette integration using NHEJ-based CRISPR-Cas9

To apply the NHEJ-based CRISPR-Cas9 method, we generated three individual plasmids (Fig. 1B). Plasmid 1 is a minicircle donor plasmid (MC-donor) harboring integration cassettes containing the Zeocin resistance gene, and the P2A self-cleaving peptide and mClover3 sequences preceded or followed by a ten amino acid linker (Glycine_4_-Serine)_2_. Thus, integration cassettes had two orientations: (Zeo)-P2A-mClover3-linker for N-terminus tagging and linker-mClover3-P2A- (Zeo) for C-terminus tagging. Plasmid 1 also contains a cleavage site complementary to a 20 bp gRNA named “Scramble gRNA” (Sc-gRNA), originally described by Suzuki *et al.* (18), followed by the PAM sequence TGG (Fig. 1B, Fig. S1). Since Sc-gRNA is not present in any protein-coding gene in the human genome, the probability of off-target activity decreases substantially with INDEL rates at off-target sites of less than 1% (18). Plasmid 2 is a CRISPR/Cas9 expression plasmid that harbors the coding sequence for SpCas9-NG under the control of a CMV promoter (20). As mentioned before, SpCas9-NG can recognize a broader range of PAM sequences which enables the design of more gRNAs per targeted integration locus of each gene to introduce in-frame DSB. Plasmid 2 also contains the Sc-gRNA that directs the Cas9 to introduce a cut into the donor DNA (Fig. 2A and Fig. S2A). Plasmid 3 contains the lenti-CRISPR gRNA library that targets all ∼20K protein-coding genes of the human genome (Fig. 2B and Fig. S2B).

**Fig. 2.**
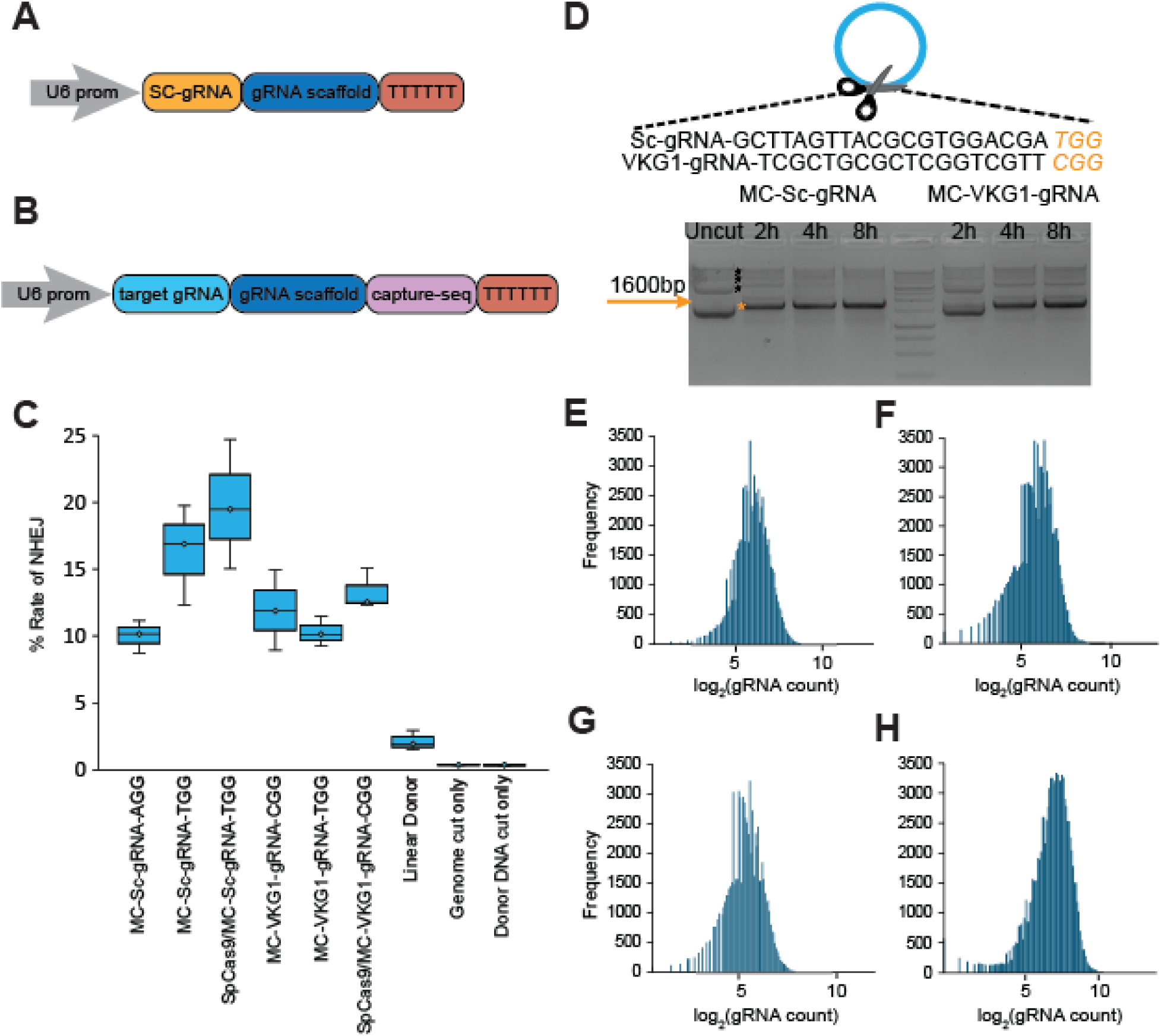
Design and optimization of seamless DNA cassette integration into the genome using NHEJ-based CRISPR-Cas9. (A) Design of donor-cleaving plasmid. The U6 promoter-based expression cassette followed by the Sc-gRNA (orange), gRNA scaffold (blue) and termination signal sequence (red). (B) Design of gRNA-CRISPR-library plasmid. The U6 promoter-based expression cassette followed by the library gRNAs targeting 18,489 human genes including their splice variants (cyan), gRNA scaffold (blue), capture sequence (pink) and termination signal sequence (red). (C) Efficiency of NHEJ-based strategies using two different gRNA cleaving vectors. The efficiency of CRISPR/Cas9-induced NHEJ-mediated DNA integration was tested by targeting five human protein-coding genes (RPS6KB1, AKT1, PKD1, RPS6 and PPP2CA) to insert four types of minicircle (MC) donor vectors targeted by distinct guide RNAs directed against different PAM sequences, TGG, AGG, or CGG (MC-Sc-gRNA-TGG, MC-Sc-gRNA-AGG, MC-VKG1-gRNA-TGG and MC-VKG1-gRNA-CGG). Box plots represent the distribution of measured values across three technical replicates for each condition. The central line within each box represents the median, while the box edges indicate the interquartile range (IQR; 25th to 75th percentile). Whiskers extend to the smallest and largest values within 1.5×IQR. We observed higher knock-in efficiency with Sc-gRNA donor using TGG as PAM (MC-Sc-gRNA-TGG) in comparison to VKG1 donors (MC-VKG1-gRNA-TGG and MC-VKG1-gRNA-CGG). Consistent with previous observations, PCR-amplified donor DNA (linear DNA) resulted in little to no knock-in events (35). We also obtained higher integration efficiency in cells that were transfected with SpCas9-NG plasmid prior to introducing donor vectors and lentiCRISPR-22-gRNAs lentivirus-infected plasmids (SpCas9-NG/MC-Sc-gRNA-TGG and SpCas9-NG/MC-VKG1-gRNA-CGG). (D) In vitro Cas9 cleavage Assay to evaluate the efficiency of our strategies for introducing double-strand breaks (DBS) into the donor vectors harboring Sc-gRNA or VKG1-gRNA target sites. The uncut DNA (lane 1) shows supercoiled conformation of minicircle DNA marked with asterisk in orange (*) and 3 possible conformations for parental plasmid marked with asterisks in black (*) (supercoiled, relaxed and nicked). Successful DSB into the donor vectors results in a band of 1600 bp in length. These results show that Cas9 efficiently introduces DSB into Sc-gRNA site within 2 hours of incubation. (E and F) gRNAs recovered after cloning designed libraries for 5’ and 3’ tagging, respectively. (G and H) gRNAs recovered after infecting HEK 293T cells with 5’ and 3’ lentiviral libraries, respectively. NGS data illustrates the close to equal distribution of gRNAs within the library.

## Optimization of seamless DNA cassette integration using NHEJ-based CRISPR-Cas9

To test and optimize our experimental design to endogenously tag protein-coding genes at both 5’- and 3’-ends, we selected a set of five human genes including RPS6KB1, AKT1, PKD1, RPS6 and PPP2CA, that had been previously tagged without detrimental consequences to their function (36). Twenty-two gRNAs were designed for these five human genes (Supplementary Table 1), and CRISPR/Cas9-NHEJ-mediated DNA integration was performed by infecting cells with a pool of the twenty-two-locus-specific gRNA-expression vectors. HEK 293T cells were co-transfected at the same time with the plasmid pX330-SpCas9-NG expressing the SpCas9-NG endonuclease and either VKG1-gRNA or Sc-gRNA guides, and one of four MC-donor vectors, targeted by guides Sc-gRNA or VKG1-gRNA followed by one of three different PAM sequences TGG, AGG, or CGG (Sc-gRNA-TGG, Sc-gRNA-AGG, VKG1-gRNA-TGG or VKG1-gRNA-CGG).

We assessed SpCas9-NG activity at the VKG1-gRNA or Sc-gRNA targeted sites. Sc-gRNA as mentioned is an artificial sequence that is not present in the human genome (18) while VKG1- gRNA is a sequence in the pUC19 plasmid backbone that has shown low off-target activity in previous studies (35). In the original studies, these sequences were recognized by wild-type Cas9 targeted by PAMs AGG and CGG, respectively. However, since we used SpCas9-NG instead of wild-type Cas9, we also tested the TGG PAM sequence because it exhibits the highest cleavage efficiency for this Cas9 variant (18, 21, 35).

Forty-eight hours after infection and co-transfection, we selected for cells successfully infected with the pool of twenty-two-locus-specific gRNA vectors, by treating them with Blasticidin (4 µg/ml) (Fig. S3A). Then, five days after infection/co-transfection, resistant cells were treated with Zeocin (100 µg/mL) for 14 days (Fig. S3B). As negative controls, we also measured the integration efficiency in cells infected/transfected with the locus-specific gRNA vectors and pX330-SpCas9- NG but no donor integration cassette; and cells only co-transfected with pX330-SpCas9-NG harboring either VKG1-gRNA or Sc-gRNA and minicircular donor plasmids, resulting in cleavage of donor plasmids but not specific gene loci. The absolute knock-in efficiencies *via* NHEJ-mediated targeted integration were measured by the ratio of live cells treated *versus* non treated with Zeocin (Fig. 2C). No knock-in was observed in the two negative controls. In the case of Sc- gRNA, we observed higher knock-in efficiency with MC-Sc-gRNA-TGG (16.34 ± 3.73%) compared to MC-Sc-gRNA-AGG (10.01 ± 1.22%). For VKG1-gRNA, we observed higher knock-in efficiency using MC-VKG1-gRNA-CGG (11.92 ± 3%) than MC-VKG1-gRNA-TGG (10.29 ± 1.1%). Notably, we observed higher knock-in efficiency with Sc-gRNA donors than with VKG1- gRNA. Our data also indicated that the knock-in efficiency was higher using TGG as PAM for Sc-gRNA. Consistent with previous observations, PCR amplified donor DNA resulted in few spurious knock-in events (2.12 ± 0.74%) (Fig. 2C) (35).

As successful gene editing requires a minimum threshold expression level of Cas9, we compared the knock-in efficiency between cells that were infected and co-transfected with the three types of editing and donor plasmids at the same time with cells that were first transfected with pX330- SpCas9-NG for 18 hours prior to infection/transfection with the twenty-two-gRNA library and donor plasmids. We observed higher knock-in efficiency when the cells were transfected with SpCas9-NG plasmid prior to introducing target specific gRNAs and donor DNA (SpCas9/MC-Sc- gRNA-TGG: 19.75 ± 4.48 %) (Fig. 2C).

We confirmed the efficiencies of MC-Sc-gRNA-TGG and MC-VKG1-gRNA-CGG-directed cleavage in cells using a biochemical assay with purified donor plasmids and Cas9 endonuclease. Consistent with our results in cells, Cas9 cleaved more efficiently MC-Sc-gRNA-TGG than MC- VKG1-gRNA-CGG donor vectors, with 90% digestion within 2-hour incubation (Fig. 2D). Collectively, we concluded that using MC-gRNA-TGG as the donor DNA and transfecting cells with pX330-SpCas9-NG prior to infection/transfection with the gRNA library and donor plasmids could be applied to the library-level integration.

## Genome-scale lentiviral gRNA library construction

To test whether our experimental design could be scaled to the library-level generation of endogenously tagged human cell lines, we targeted the entire human protein coding genes for donor cassette integration. We used the following criteria in designing a genome-wide gRNA library: **i**, gRNAs only introduce double strand breaks in frame; **ii**, minimal off-target activity; **iii**, maximal on-target activity; **iv**, avoidance of homopolymer stretches (e.g., AAAA, GGGG) (37); and **v**, GC content between 20 to 80%. This resulted in two gRNA libraries consisting of 73,909 and 85,394 unique guides designed for 5’- and 3’-end tagging, respectively (Supplementary Tables 2 and 3). These libraries contain a minimum of 1 and a maximum of 8 gRNAs per splice variant of human genes (Fig. S4 A and B). gRNAs were synthesized as 79-mer oligonucleotides on microarray chips and amplified by PCR as a pool. The PCR products were cloned into a modified version of lentiviral vector ‘lentiCRISPR-V2’ to yield at least 2000-fold representation per gRNA in each library. We evaluated the diversity and coverage of the library using next-generation sequencing (NGS) (Supplementary Tables 4 and 5). This analysis showed 99.8% and 99.9% coverage of the designed sgRNAs for each N and C library. In addition, a skew ratio below 5 indicated uniform representation, minimizing bias and optimizing screening efficiency. Guide representation ranged from 5% to 95%, ensuring proportional distribution across the library, and maximum screening efficiency and identification of a hit (Fig. 2E and F). A similar result was observed for the diversity and coverage of the gRNA libraries after infection of HEK 293T cells, illustrating the efficiency of the lentiviral particles to deliver the ∼ 80,000 gRNAs into the genomes of these cells (Fig. 2G and H) (Supplementary Tables 6 and 7).

## *In vivo* genome-wide editing and fusion of reporter sequences to protein-coding genes

To test PRISM across the 18,489 protein coding genes, we tested for integration of the coding sequence for the green fluorescent protein variant mClover3 fused to Zeo in HEK 293T cell line. Similar to our small-scale test above, the donor DNA had two orientations, linker-mClover3-P2A- Zeo and Zeo-P2A-mClover3-linker (Fig. S5). With these cassettes expressed in-frame with the protein-coding sequences, we could select for donor integration with Zeo and for functional integration by visualization of mClover3 fluorescence. HEK 293T cells were transfected with pX330-SpCas9-NG/Sc-gRNA and 18 hours later, cells were infected with 5’ or 3’ lentiviral gRNA libraries at a multiplicity of infection (MOI) of 0.3 and transfected with minicircle plasmid encoding the donor cassette.

To select for successful infection of library gRNAs, 48 hours post transduction cells were treated with 4 µg/ml Blasticidin (Fig. S3A). Five days later, to select for successful integration of donor cassette, cells were treated with 100 µg/mL Zeocin (Fig. S3B) until all the non-transduced cells in the negative control died out. Afterwards, cells were kept in culture for 21 days, passaging every 4 days. To avoid bottlenecking, we aimed to keep the total cell count above 3.7 x 10^7^ and 4.3 x 10^7^ for 5’- and 3’-end tagging, respectively, corresponding to a 500-fold coverage of library diversity. Successful integration was evaluated by screening cells for mClover3 fluorescence. We observed mClover3 signal across mixed populations of selected cells with heterogeneous expression levels inferred from the intensity of fluorescence and unique subcellular localizations, suggesting successful functional tagging of a wide variety of genes (Fig. 3A).

**Fig. 3:**
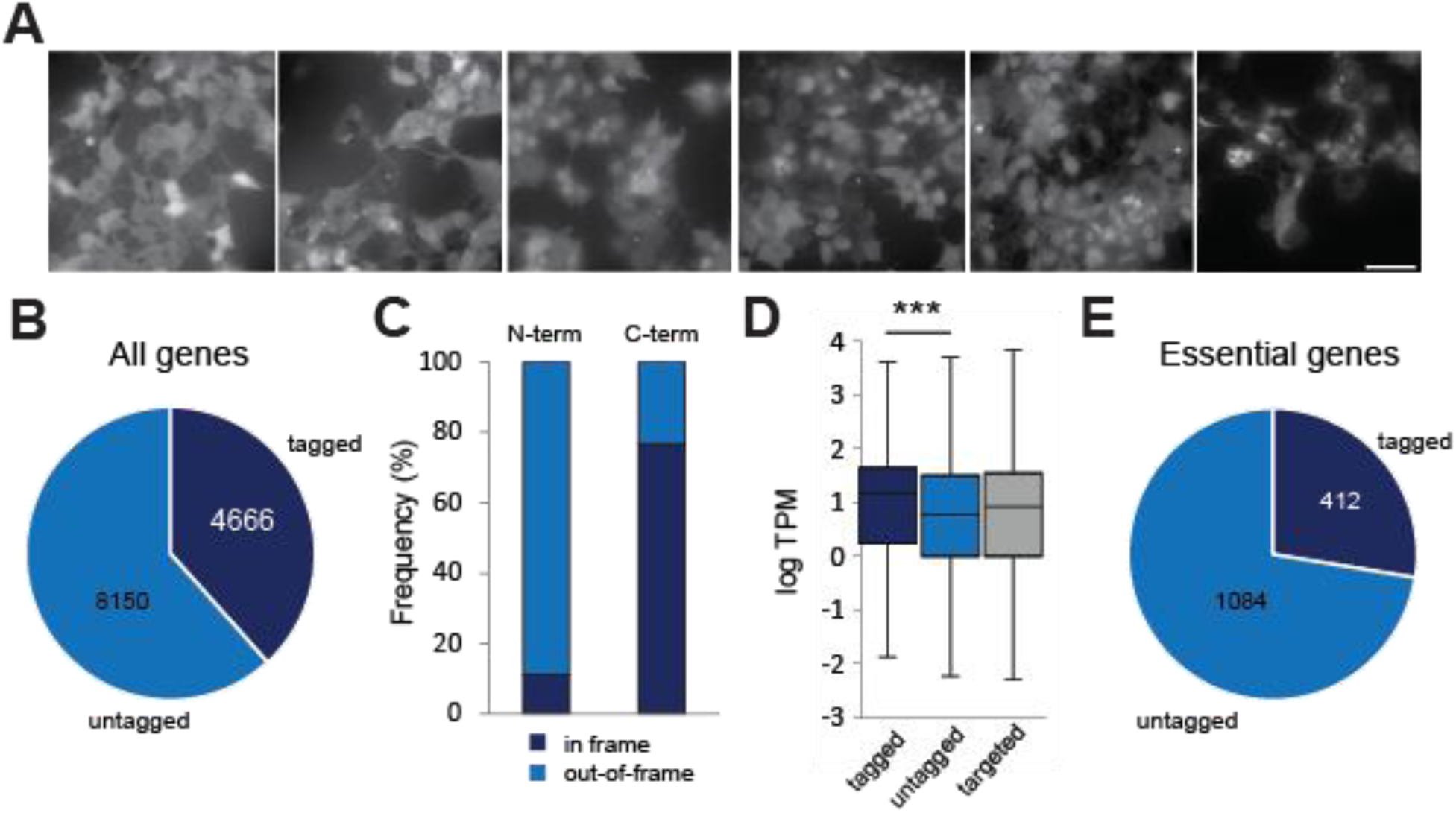
Genome-wide CRISPR integration of reporter protein donor DNA fragments. (A) Mixed pools of cells in which endogenous proteins are fused to mClover3 by PRISM. Cells vary in fluorescence intensity and patterns of expression suggesting varying abundances and subcellular localizations of the tagged proteins. Scale bar is 50 µm. (B) NGS results indicate that ∼4666 of the targeted genes were tagged 5’ or 3’ to protein-coding sequences in the genome of HEK 293T cells. (C) Frequency of in-frame/out-of-frame events at the integration sites for 5’ or 3’ libraries containing the tag and the target coding sequence. (D) Distribution of abundance for targeted proteins expressed in HEK 293T versus tagged or untagged targets. Boxes represent 25th and 75th percentile, and whiskers represent minimum and maximum values. Median is indicated as middle line and statistical significance was calculated by Student’s t-test. Data indicates that low protein abundance constitutes a significant challenge to successful tagging in HEK 293T cells. (E) NGS results indicate that ∼30% of the essential genes were tagged 5’ or 3’ to protein-coding sequences in the genome.

Once the screen was completed, a fraction of the integrated library was used to identify correctly tagged genes for each site (5’ or 3’). Briefly, genomic DNA was extracted from one sample of the cells that survived Zeocin selection followed by physical DNA fragmentation. Next, a poly-C tail was added to the 3’ ends of the fragmented DNA by using a terminal deoxynucleotidyl transferase (TdT). Regions containing the junctions of the donor integration sites were enriched for by a first round of PCR with primers that recognized the tag and the C-tail. Next, a second round of PCR was performed to insert adaptors needed for sequencing (38). Amplified products were subjected to high-throughput sequencing using Illumina NextSeq (see methods). The correct paired reads with forward and reverse reads mapped within the defined DNA fragment length on the human genome were then sorted; and the read pairs overlapping with genomic locations targeted by gRNAs in the library were retrieved. These analyses showed that a total of 4,666 genes out of the 18,489 targeted were correctly tagged in this study (Fig. 3B), including 3,037 at the 5’ and 2,031 at the 3’ (Supplementary Tables 8 and 9). As expected, a higher number of genes were tagged in 5’ *versus* 3’ given that the antibiotic expression for the 5’ cassette does not depend on in-frame integration. Since we could only process a fraction of the tagged libraries to directly identify integration sites, due to the high coverage of the library and the limited amount of DNA that could be processed per step using the mentioned approach, it is possible that more genes were correctly tagged but not detected. Regardless, and considering that around 12,816 genes are reported to be expressed in HEK 293T cells, approximately 36% of expressed genes in this cell line were tagged (39) (Fig. 3B). This represents the highest number of tagged protein-coding genes that has been reported for a single pooled study to this day, and a ninefold increase over what was previously reported (11).

Analysis of integration site sequences showed that for the 3’ tagged library, 76.7% of the insertions were in-frame between the targeted gene and the donor cassette while for the 5’ library only 11.2% were in-frame (Fig. 3C). We expected the number of out-of-frame events to be higher for the 5’ library given that the reading frame-independent expression of the antibiotic resistance gene can occur. However, some of the out-of-frame polypeptides produced could also be toxic to the cells, resulting in further selection against out of frame fusions. For codon deletion/insertion at the integration sites, we observed a broad distribution with higher frequency of deletions/insertions closer to the integration sites (Fig. S6).

We then assessed whether gene intrinsic expression levels could have influenced the donor cassette integration efficiency of the targeted genes, since at least for the 3’ insertion, the survival of the cells carrying correctly tagged genes depends on the strength of the specific promoter for the targeted gene. Using RNA-seq measurements performed in HEK 293T cells (8) as proxy for protein abundance, showed that on average, tagged genes have higher expression levels than untagged genes, represented as log-transformed Transcripts Per Million (log TPM) (Fig. 3D). Thus, expression level was a variable that affected endogenous tagging, as it has been seen before for other high throughput donor cassette integration strategies (8, 11).

We next analyzed the efficiency of our strategy for tagging essential genes, which are often the target of most scrutiny for functional characterization studies but can also be most sensitive to manipulation. PRISM should work as a filter, selecting for N- or C-terminal fusions of proteins to essential gene products that do not result in functional disruption, due to destabilization, degradation or mis-localization of the protein. For this analysis, gene essentiality annotations were obtained from the Cancer Dependency Map (DepMap) 25Q3 release (40). We found that 412 essential genes were tagged at both 5’- and 3’-ends (N- and C-terminal tagged), 249 uniquely at 5’ (N-terminal tagged) and 195 only at 3’ ends (C-terminal tagged) (Fig. 3E). This amounts to a 27.5% success rate. These results illustrate a key application of PRISM, creating a roadmap for the seamless and systematic integration of tags into protein coding genes including essential genes, to generate arrays of cell lines with specifically labelled proteins. Thus, PRISM is a screen that allows for selection of the best sgRNAs and preferable N- or C-terminal tagging orientation to minimally disturb the function of a protein.

Finally, we assessed PRISM capability in other cell lines. To test this, we performed PRISM in three additional human cell lines: HeLa, colorectal carcinoma HCT116, and near haploid myelogenous leukemia Hap1. For practical considerations, while we performed CRISPR-NHEJ knock-in with 500-fold coverage in HEK 293T cells, we opted for a lower 300-fold coverage in these other cell lines as a simplified proof-of-concept. We also used a faster and simpler approach to estimate the number of genes that could have been tagged by amplifying and sequencing the regions corresponding to the locus-specific-gRNA/gRNA-scaffold present in the pool of genomic DNA after antibiotic selection. After the screen was completed, genomic DNA was extracted from one sample of the cells, and gRNA regions were amplified followed by deep sequencing. NGS data were analyzed to identify and quantify the specific gRNAs coding sequences present in the pool of genomic DNA by mapping reads to the entire designed oligo pool. Our results showed the presence of a diverse pool of gRNAs across all three cell lines, including those targeting 1565 genes in HCT116, 1360 genes in HeLa, and 11,668 genes in Hap1 cells (Fig. S7 and Supplementary Table 10). Further tests with optimized parameters for each cell line including gRNA libraries and infection/transfection conditions will be required to better compare PRISM efficiency between different cell types.

## Cellular localization of fluorescently tagged proteins for PRISM single clones

We next evaluated PRISM performance at the level of single integration clones. To achieve this, we randomly selected 96 putative single clones from our mClover3-pooled library, which we expanded and imaged individually. Out of these, 76 clones displayed detectable signal over background (80%). However, sequencing analysis to identify the target protein showed mixed populations for 24 samples. The remaining 52 clones were analyzed by microscopy, and our localization annotations were compared to those reported in the Human Protein Atlas (HPA), which is a reference immunohistofluorescence-based collection of human subcellular protein localizations (41). This analysis showed that localization annotations matched for 92% of the proteins tagged (Fig. 4 A and B). It is important to note, however, that HPA mainly reports on a wide variety of cells aside from HEK 293T, so we did not expect a perfect overlap since proteins can vary in their localization across related compartments in different cell types (Fig. 4B and Fig. S8).

**Fig. 4:**
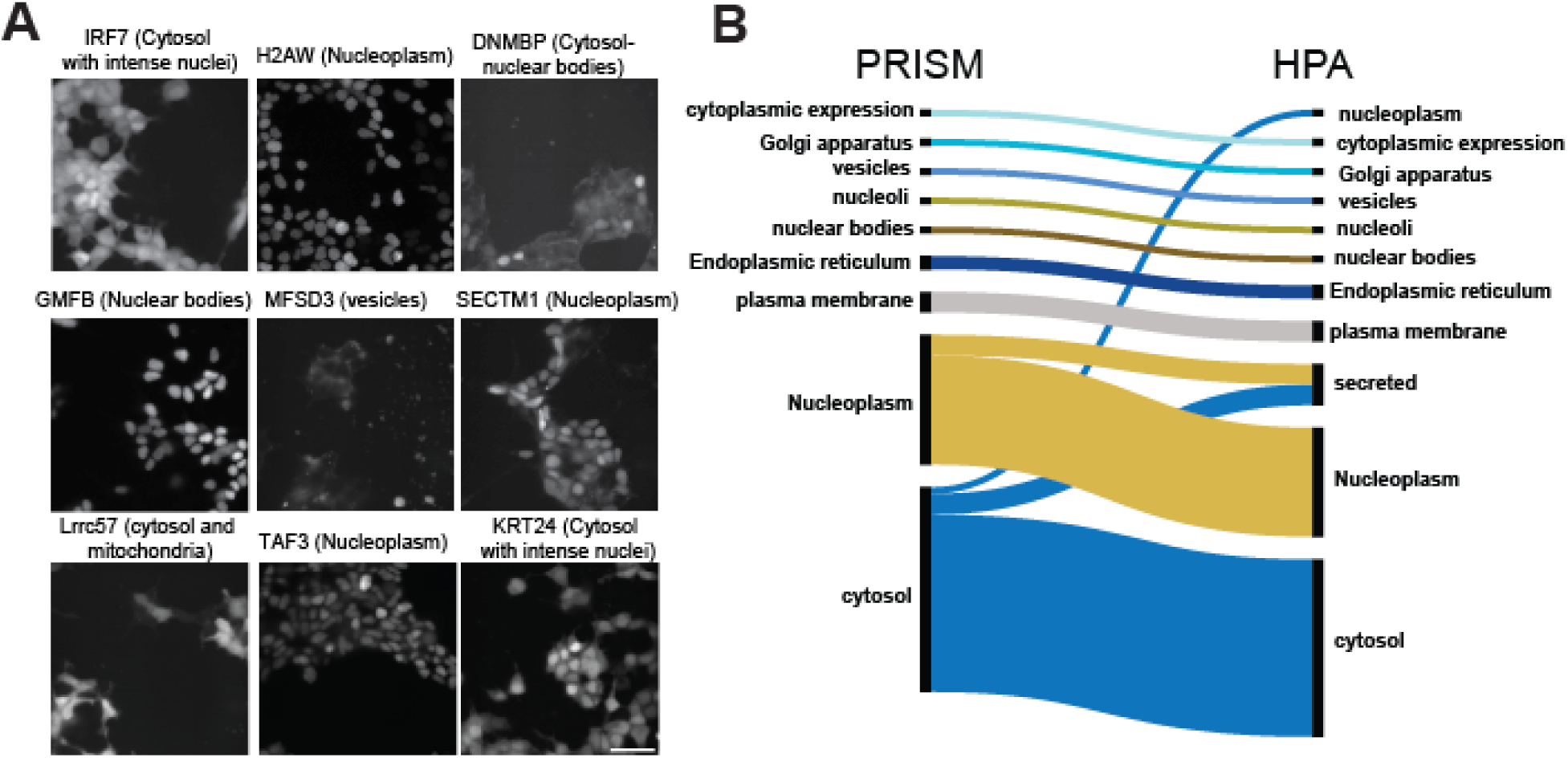
Fluorescent analysis for single clones of CRISPR-NHEJ tagged coding sequences. (A) Imaging analysis of individual successful target proteins. mClover3 fluorescence intensity and subcellular localization vary widely for each gene. (B) Comparison of annotated protein localizations between PRISM and the Human Protein Atlas (HPA) datasets (41). The diagram features colored bands that correspond to groups of proteins with similar localization annotations between our dataset and HPA. The width of each band is proportional to the number of proteins in the group.

## Discussion

Here, we report a robust approach for the rapid and efficient generation of reporter protein-tagged endogenous human gene sequences. Critical to the efficiency of PRISM was the use of circularized mini plasmids as donor vectors and their intracellular cleavage. Plasmids have a longer lifetime than linear DNA, and proteins involved in NHEJ, such as KU70 and KU80, can attach to the 5′- or 3′-ends of donor DNA in response to DSB in cells, which protect donor DNA from degradation and facilitate ligation of the donor to genomic DNA (42). Compared with existing methods, PRISM has the following advantages: First, in contrast to multistep preparation of HDR targeting vectors, we designed universal donor cassettes for all ∼20,000 human proteins that can be generated rapidly and in large scale without any requirement for target sequence-specific donor vector reconstruction. Additionally, the delivery of the donor cassette as minicircle plasmids, aside from increasing its lifetime, minimizes undesired integration of backbone plasmid leading to higher donor integration efficiency. Seamless HDR tagging of protein genes will remain important for systematic, quantitative spatiotemporal analysis of proteins in living cells, and measuring protein abundances and protein-protein interactions(7, 43). However, PRISM will aid in screening for functional fusions of genes that have proven recalcitrant to tagging with reporters and to obtaining endogenous fusion of many more genes that are currently unavailable including in different cell lines. Second, unique to PRISM is the use of SpCas9-NG having an expanded number of PAM recognition sites including some considered inaccessible or non-targetable with other Cas9 variants (19, 21). Studies analyzing the off-target activity of Cas9-NG have shown either similar values with different profiles compared to SpCas9 (21); increased off-target numbers (44); or higher mismatch tolerance, which could imply possible higher off-target events (45). Even if Cas9- NG had increased off-target activity, this could be balanced with more stringent selection of the sgRNAs, since off-target activity is mainly guide-dependent (46). Third, also distinctive to PRISM, is the design of a library with multiple sgRNA per gene. SgRNA efficiencies can vary greatly and selecting the optimal is not straightforward, so having more than one sgRNA per gene increases the chances of editing at selected genomic sites. Another advantage of the PRISM sgRNA library design, seen in other studies, is that of selecting in-frame guides or guides with the same reading frame once Cas9 cuts the targeted sequence (11, 14). Fourth, also related to the donor cassette, the P2A-antibiotic donor design in our approach provides a simple solution to select only cells with in-frame 3’ donor DNA and decreases the errors introduced by NHEJ. In the case of out-of-frame 5’ integrations, these will be discarded during secondary selection or measurements performed on cells expressing the fusion proteins. Finally, the utility of endogenously tagged proteins enables the function of a protein to be characterized under the control of native regulators of gene expression and without disturbing endogenous interaction stoichiometry.

We envision that PRISM could be used to manipulate protein abundances or interactions in a variety of ways, including chemically induced degradation or proximity, such as auxin-induced single or pairwise protein degradation or mandipropamid-induced forced proximity of protein pairs (47–50). Similarly, measuring protein abundances or protein-protein interactions could be achieved with protein-fragment complementation assays, such as those based on dihydrofolate reductase reporter (51, 52) or split fluorescent proteins (43, 53).

There are several ways PRISM could be further optimized depending on the intended application. In its current form, PRISM employs the same Cas9 protein to cleave both the donor mini-plasmid and the genomic target site (Fig. 1). This dual use introduces competition between the target-specific gRNA and the donor-specific gRNA for Cas9 binding, which may reduce cleavage efficiency at either site, ultimately lowering insertion efficiency (54, 55). One potential solution, as demonstrated with HITAG (11), is to use two orthogonal Cas9 variants—one for the donor and another for the target site in the genome—to avoid this competition. Additionally, for medium-scale libraries (rather than proteome-wide screens), higher-fidelity Cas9 variants, such as xCas9 (45), can be employed to reduce off-target effects. However, this comes at the cost of a more restricted range of PAM sites, cleavage efficiency (21), and hence applicable for medium scale libraries. Additional optimization of PRISM could be inclusion of tandem copies of the selection marker in the donor integration cassette. Since the expression level of the resistance gene fused to the desired tag depends on the strength of the targeted gene promoter, this affects the donor integration success rate for lowly expressed genes. Adding more copies of the selection enzyme gene to the donor cassettes have proven to increase the number of viable cells after antibiotic selection (11). Other variables that could enhance the performance of PRISM are the optimization of the transfection conditions, including amounts and ratios of Cas9- and donor-coding plasmids added to the cells. Donor integration efficiencies can also be improved by selecting sgRNAs as close (less than 10 bps) as possible to insertion sites (11, 56). Here, the expanded PAM recognition repertoire of Cas9-NG is an advantage, because it is more likely that a PAM closer to the insertion site exists. PRISM donor insertion efficiency could also be increased by combining several of these strategies, however, without all these optimizations, the number of genes tagged in this study is considerable despite targeting a genome wide library which imposes many more challenges than smaller libraries.

Altogether, PRISM provides universality, scalability, specificity, selectability and paves the way for the genome-scale construction of human cell lines tagged with different functional tags at endogenous loci. Consequently, PRISM makes whole proteome tagging and functional genomics readily accessible to any researcher aiming to manipulate or measure the abundances or interactions of proteins.

## Methods

### Plasmid construction

#### Constructing plasmid harboring SpCas9-NG and donor cleaving gRNA

The plasmid coding Cas9-NG (21) was purchased from Addgene (pX330-SpCas9-NG, Plasmid #117919)) and contains two expression cassettes, a human optimized SpCas9-NG and a single guide RNA. The guide RNA expression cassette was used to individually clone two donor-cleaving gRNAs named “scramble-gRNA” (Sc-gRNA) GCTTAGTTACGCGTGGACGA (18) and VKG1- gRNA TCGCTGCGCTCGGTCGTT(35). The gRNA sequences were synthesized as single stranded DNA primers (Supplementary Table 11), amplified by PCR and cloned 3’ to a U6 promoter following the protocol of Ran *et al.* (57). Briefly, gRNA forward and reverse primers were annealed at 100°C in a mixture of equal molar concentrations and volumes and gradually cooled down to room temperature. The annealed sequences were then used as inserts and ligated by T4 DNA ligase (New England Biolabs) into the pX330-SpCas9-NG plasmid previously digested by BbsI (New England Biolabs). The ligation products were transformed into Stbl3- competent cells and the resultant clones were screened with restriction enzymes and verified by sequencing. The Sc-gRNA was artificially designed and does not exist in human, mouse or rat genomes, and is reported to successfully guide the introduction of double-strand breaks into a donor DNA flanked with complementary sequences using wild-type Cas9 (18, 21). The VKG1- gRNA sequence is part of the pUC19 vector backbone and is reported to have few off-target sites in human and mouse genomes(35).

#### Constructing donor vector for 5’ and 3’ tagging of protein-coding gene sequences

To make donor vectors Zeocin-P2A-mClover3-linker and linker-mClover3-P2A-Zeocin for tagging human genes at respective 5’ and 3’ ends, the coding sequences for the Zeocin resistance gene Sh ble and the P2A peptide were PCR amplified from the LentiCRISPR-v2 plasmid (Addgene, Plasmid #52961) (Supplementary Table 11). The coding sequence for the mClover3 reporter was amplified from the pMK290 (mAID-mClover-Hygro) plasmid (Addgene, Plasmid #72828). All the fragments were assembled *in vitro* using Gibson assembly (New England Biolabs) and were flanked with complementary sequences of Sc-gRNA or VKG1-gRNA followed by either AGG or TGG as PAMs for the Sc-gRNA, and CGG or TGG for VKG1-gRNA (Supplementary Table 11). The tested and reported PAMs for Sc-gRNA and VKG1-gRNA are AGG and CGG, respectively. Both were tested using wild type Cas9. We also tested TGG as PAM for these two gRNAs because the SpCas9-NG variant we used, has been reported to have highest cleavage efficiency at TGG sequences (21). In the next step, the fragments were cloned into ApaI and SmaI sites of the minicircle producer plasmid pTubb3-MC (Addgene, Plasmid #87112) using GeneArt seamless cloning (Thermo Fisher Scientific). In this design, the expression of the Zeocin resistance marker and mClover3 genes are driven by the endogenous promoter of the protein coding genes to which the mClover3-coding oligonucleotide donor is fused.

#### Construction of Lenti-CRISPR-gRNA library

We used a modified version of LentiCRISPR-V2 (58) plasmid for cloning the gRNA libraries. In this plasmid, we first replaced the sequence coding for Cas9 with that for mCherry using PCR followed by GeneArt seamless cloning (Supplementary Table 11). This plasmid also contains an expression cassette for gRNAs including the gRNA scaffold sequence (Fig. S2B). We then added a short sequence named “Capture Sequence” (GCTTTAAGGCCGGTCCTAGCAA) (59) immediately before the gRNA termination signal, which elongates the 3’ end of the gRNA (Fig. S2B, Supplementary Table 12). The capture sequence is a 22 bp DNA fragment, designed and used in feature barcoding technology to directly capture gRNAs present in a cell inside a gel-bead-in-emulsion (59). Dixit *et al.* showed that this short fragment does not affect the gRNA scaffold. We used this sequence to provide a priming site to later amplify the gRNA library in individual cells and to help distinguish the donor-cleaving gRNA from the gRNA library in later steps (59) since both gRNA cassettes contain the same promoter and scaffold sequences (Fig. S2). gRNA scaffold followed by the Capture Sequence was synthesized as a single stranded DNA (IDT) (Supplementary Table 12) and was cloned into the LentiCRISPR-V2 plasmid using GeneArt seamless cloning kit (Thermo Fisher Scientific).

#### Minicircle DNA vectors preparation

Production of minicircle DNA vectors (MC) were performed in accordance with the manufacturer’s protocol (System Biosciences (SBI)) with some modifications. Briefly, the final minicircle constructs, pre-minicircle plasmids P2A-Zeocin-MC, Zeocin-P2A-MC and linker- mClover3-P2A-Zeocin-MC with Sc-gRNA targeting sequences were introduced into the ZYCY10P3S2T *E. coli* minicircle producer competent cells (SBI) and amplified overnight in Terrific Broth (TB) (pH 7.0) (Fisher Scientific). The minicircle production was induced by mixing the overnight TB culture with an equal volume of minicircle induction mix comprising fresh LB and 20% L-arabinose, followed by a 5 h incubation at 30°C with shaking at 250 rpm and 2 h incubation at 37°C with shaking at 250 rpm. Minicircle DNA was then isolated with ZymoPURE II Plasmid Maxiprep Kit (Zymo Research). Finally, 1.0 μg of each minicircle plasmid (P2A- Zeocin-MC, Zeocin-P2A-MC and linker-mClover3-P2A-Zeocin-MC) was digested with SmaI restriction enzyme (New England Biolabs) and then run on a 1.5% agarose gel to determine plasmid quality (Fig. S9).

#### *In vitro* Cas9 cleavage assay for donor cleaving gRNAs efficiency

We checked the cleavage efficiency of both Sc-gRNA and VKG1-gRNA toward donor DNA using an *in vitro* Cas9 Cleavage Assay. To prepare each donor-cleaving gRNA for *in vitro* transcription, we followed methods by Lin *et al.* with some modifications (60). gRNAs were obtained by *in vitro* transcription of a DNA template of the following sequence: 5′-TAA TAC GAC TCA CTA TAG GNNNNNNNNNNNNNNNNNNNG TTT AAG AGC TAT GCT GGA AAC AGC ATA GCA AGT TTA AAT AAG GCT AGT CCG TTA TCA ACT TGA AAA AGT GGC ACC GAG TCG GTG CTT TTT TT-3′, containing a T7 promoter (TAATACGACTCACTATAG), ∼20-nt GCTTAGTTACGCGTGGACGA for transcription of Sc-gRNA or 18 bp sequences of VKG1- gRNA (TCGCTGCGCTCGGTCGTT) followed by gRNA scaffold. The DNA templates were generated individually *in vitro* using Gibson assembly (New England Biolabs) (Supplementary Table 12). The resulting dsDNA was then purified using NucleoSpin Gel and PCR Clean-Up (Takara) and approximately 200 ng of DNA was used as template for a T7 *in vitro* transcription (IVT) reaction (MAXIscript T7 Transcription Kit from Invitrogen) generating the gRNAs. In the next step, *in vitro* transcribed gRNAs were treated with DNase (DNaseI-Thermo) and purified using the Omega EZNA PF kit. The concentrations of the purified gRNAs were then measured using a NanoDrop. After purification, each gRNA was diluted to 200 ng/µL and stored at –80°C. Cleavage assays were conducted in a reaction volume of 20 μL containing SpCas9 (homemade) (200 nM), each gRNA (400 ng), and each donor plasmid (1000 ng) in 1X cleavage assay buffer (homemade) at 37°C, tested over the time periods of 2, 4 and 8 hours. The cleaved dsDNA was analyzed by 1.5% agarose gel electrophoresis (Fig. 2D).

#### Cell culture

HEK 293T and HeLa cells were cultured in Dulbecco’s modified Eagle’s Medium (DMEM, Wisent) supplemented with 10% fetal bovine serum (FBS, Wisent) and 1% Penicillin-Streptomycin (Pen/Strep). Hap1 cells were cultured in Iscove’s Modified Dulbecco’s Medium (IMDM, Wisent), 10% FBS and 1% Pen/Strep. HCT116 cells were cultured in McCoy’s 5A Medium (Wisent), 10% FBS and 1% Pen/Strep. All cell lines were maintained at 37°C in a 5% CO_2_ incubator.

#### Dose-response curve for antibiotic selection of mammalian cells

We measured the kill curve for each of the antibiotics to determine the minimum amount of the antibiotic required to eliminate non-infected or non-edited cells over the course of 7 days. This was accomplished by performing the assay using cell counting kit-8 (SIGMA). HEK 293T cells were seeded in 24-well plates 24 hours prior to antibiotic treatment. Cells were treated with Zeocin (Gibco) at concentrations ranging between 0 (negative control) and 500 μg/ml for 7 days. For Blasticidin S HCl (Thermo Fisher Scientific), cells were treated with concentrations between 0 (negative control) and 15 μg/ml for 7 days. At the end of the assay, cells were treated with Cell Counting Kit-8 (CCK-8, Sigma) according to manufacturer’s instruction and an estimate of the number of surviving cells was made based on OD_450_ measurements (fig. S3A and B).

#### Design and optimization of donor DNA knock-in based on NHEJ DNA repair

We aimed to develop an efficient method for inserting exogenous DNA fragments into the human genome by harnessing the endogenous non-homologous end joining (NHEJ) DNA repair pathway. NHEJ is the most efficient repair system in mammalian cells since it is functional during all phases of the cell cycle(61). Our design consists of two main steps: First, simultaneous introduction of double-strand breaks (DSB) in both genomic and donor vectors inside living cells and second, integration of the linearized donor vector into the cleaved genome by NHEJ DNA repair system (Fig. 1A and B). To efficiently apply the NHEJ-based method, we optimized three individual plasmid elements: i, our modified version of the SpCas9-NG plasmid (derived from pX330) that harbors Sc-gRNA or VKG1-gRNA, ii, the mini-circular donor vectors harboring coding sequences for the Zeocin resistance gene (Zeo) fused to a P2A self-cleavage peptide and flanked with complementary sequences of Sc-gRNA or VKG1-gRNA followed by either AGG or TGG as PAMs for the Sc-gRNA, and CGG or TGG for VKG1-gRNA and iii, our modified version of the lentiCRISPR-V2-vector harboring locus specific gRNAs. To evaluate the efficiency of our NHEJ method for integration of donor DNA into the genome, we first targeted five human-specific genes including RPS6KB1, AKT1, PKD1, RPS6 and PPP2CA. These genes were chosen since they have diverse expression levels, which could be an ideal representation of the broad distribution of transcript levels for the human genome. Additionally, we have previously tagged these genes without functional consequences(62). We made a lentiCRISPR-gRNA library consisting of twenty-two locus specific gRNAs targeting above-mentioned genes using protocol established by Ran *et al.* (57) (Supplementary Table 1).

To test for integration efficiency, well-dissociated HEK 293T cells were seeded on 6-well plates (Corning) 24 h before transfection, at a density of 2 × 10^6^ cells per well in a total volume of 2 ml of complete DMEM medium (DMEM, 10% FBS and 1X Sodium Pyruvate). Cells were transfected with a mixture of 3 µg plasmid coding SpCas9-NG and 3 µg of each donor-plasmid pre-mixed with X-tremeGene 9 transfection reagent (36 µl) (Roche) according to the manufacturer’s protocol; and infected at the same time with 200 µl of Lentiviral lentiCRISPR-22-gRNAs. As successful gene editing requires a minimum threshold expression level of Cas9, we also tested the integration efficiency of our model in cells that were transfected with SpCas9-NG plasmid 18 hours before introducing donor vectors and lentiCRISPR-22-gRNAs. To select for transduced cells, in the next step, cells were treated with 4 µg/ml of Blasticidin S over 7 days. Five days after transduction, cells were selected following Zeocin treatment (100 µg/mL) until all the non-transduced cells and cells without successful integration were dead. At the end of the screening, cell viability was measured using Cell Counting Kit-8 (CCK-8, Sigma) and used to evaluate the percentage knock-in efficiency. Results were obtained from three replicate wells and presented as average efficiency ± standard error.

#### Designing two custom gRNA libraries for N- and C-termini tagging protein-coding sequences

For the selection of coding regions using the UCSC Table Browser (63), we downloaded the table “RefSeq Curated” from human assembly hg38 (accessed on 27/03/20), group “Genes and Gene Predictions” and track “NCBI RefSeq”. We excluded non-coding entries from the table by filtering out rows with “NR_” accession prefixes. We also downloaded the “ccdsInfo” table (hg38 –> Genes and Gene Predictions -> CCDS). We created a new table by merging both tables on RefSeq accessions, thus linking protein-coding transcripts (NM) to their CCDS entry. Finally, we removed entries belonging to chr_*alt and chr_*fix. The final table contains 18,489 unique gene IDs and 38,788 unique RefSeq accessions (Supplementary Table 13). For the design of guide RNA libraries, we used the command-line version of CHOPCHOPv3(63) to find guides for our targets. We forked the source code from Github and implemented the following in-house features: Limiting the guide search to either of the terminal exons, detecting the frame of candidate guides and computing the number of residues removed from the original reading frame following non-homologous knock-in. These additions, combined with the removal of several features from the original program, allowed us to minimize the search space by pre-emptively excluding guides that failed to meet the criteria for successful knock-in. The search was limited to guides of length 20 nucleotides with the protospacer-adjacent motifs (PAMs) NGN, NAH and NTG. For each accession in our master table, we used the program to find guides that cut in-frame and trimmed at most three residues from either terminus. We further constrained guides to have a minimum GC content of 20% and a maximum of 85%. Guides with oligo(dT) tracts of four or more were excluded to avoid premature transcription termination by RNA Pol III(64)^.^ Guides with restriction motifs “ACCGGT”, “GGTACC”, “GAATGC” and “CGTCTC” were also excluded. These steps resulted in lists of 89,641 and 85,942 unique guides for 5’ and 3’ (N- and C-terminal protein fusions), respectively. Then, for the exclusion of guides based on target features, we used the “Retrieve/ID Mapping” tool of the UniProt(65) database (accessed on 20/05/20) with our list of RefSeq accessions to obtain the corresponding UniProt entries. We kept the reviewed entries only and queried them for “Molecule processing” features using the Proteins API (66)^.^ Specifically, we recorded signal peptide, transit peptide and pro-peptide annotations found at either terminal in a table (Supplementary Table 14). Using this table, we excluded all guides targeting terminal regions corresponding to either of the sequence features above. Finally, we removed duplicated guides to obtain 73,909 and 85,394 unique guides for N- and C-termini, respectively (Supplementary Table 2 and 3). Thirty-four genes had no guides passing these filters (Supplementary Table 15).

#### Genome-scale lentiviral gRNA construction

All 79-mer oligonucleotides on corresponding microarray chips for N- and C-terminal libraries (Custom Array) were amplified by PCR as a pool (Supplementary Table 11). The PCR products were then purified using NucleoSpin Gel and PCR Clean-Up (Takara) and cloned into a modified version of the lentiviral vector LentiCRISPR-V2 (Addgene Plasmid #52961) containing the mCherry gene. For this, the LentiCRISPR-V2 vector was digested with BsmBI-V2 (New England Biolabs) for 1 hour at 55°C and gel purified using the NucleoSpin Gel and PCR Clean-Up (Takara). In the next step, the purified library was sub-cloned into the digested lentiCRIPSRv2 vector at a ratio of 1:10 vector-to-insert molar ratio, using a Gibson assembly kit (New England Biolabs). The ligation reactions were precipitated using isopropanol and transformed into Endura Electrocompetent cells (Lucigen). To yield a 2000-fold representation of each library, 40 identical ligation reactions were pooled and purified, followed by 48 parallel transformations. Outgrowth media from transformations were pooled and plated onto fifty 500 cm^2^ LB-carbenicillin (100 mg/ml) agar plates and were incubated at 37°C for 12 hours. Colonies were scraped off the plates, pooled, and the plasmid DNA was extracted using the ZymoPURE II Plasmid Maxiprep Kit (Zymo Research). The extracted plasmids for each library were subjected to PCR reactions using NEBNext high-fidelity polymerase (New England Biolabs) (Supplementary Table 16) and cleaned up using the Zymo-Spin V with Reservoir. NGS was performed on purified samples on Nextseq 150 cycles, Mid Output, v2.5 (Illumina). We aimed for at least 100 reads per unique gRNA.

#### Counting gRNAs from NGS data

To count sgRNAs from genomic DNA, fastq files were analysed using MAGeCK (version 0.5.9.4) using default parameters (67).

#### Lentivirus production

gRNA library lentiviruses were produced by co-transfection of lentiviral vectors pCMV-Delta 8.9 plasmid (packaging vector), pCMV-VSV-G (envelope vector) and lentiCRISPR-V2 library plasmid, using X-tremeGene 9 transfection reagent (Roche). Briefly, the well-dissociated HEK 293T cells were seeded in T225 flasks 20–24 h before transfection at a density of 2 × 10^7^ cells per flask in a total volume of 45 ml complete medium. Overall, four flasks were used for each library. On the day of transfection, cells with 80% confluency were transfected with a mixture of pCMV-Delta 8.9 plasmid (9 µg), pCMV-VSV-G (7.7 µg), lentiCRISPR-V2 plasmid library (12 µg), and X-tremeGene 9 (270 µl), according to the manufacturer’s protocol. At 18 hours after transfection, the medium was replaced with pre-warmed complete DMEM medium. The expression level of mCherry was used as the control for estimating the transfection efficiency. The first virus-containing medium was harvested two days after transfection; cellular debris was filtered out using Millipore’s 0.45 µm Stericup filter unit and stored at -80°C. To obtain sufficient library coverage, virus particles were collected from four 225 cm^2^ flasks for each library.

#### Pooled genome-wide CRISPR knock-in in HEK 293T for integration of exogenous donor DNA fragments

To expand PRISM to the integration of linker-mClover-P2A-Zeocin and Zeocin-P2A-mClover- linker cassettes to the entire human protein-coding genes, we harnessed NHEJ DNA repair on a large scale as follows: A total of four 225-cm^2^ flasks each containing 2 × 10^7^ HEK 293T cells were seeded 20 hours prior to transfection for 5’ and 3’ integration. On the day of transfection, each flask was transfected with donor cleaving vector (SpCas9-NG-Sc-gRNA). At 18 hours after transfection, cells were simultaneously infected with lentiCRISPR-gRNA library and mini-circular plasmid harboring linker-mClover-P2A-Zeocin, or Zeocin-P2A-mClover-linker donor vectors. The infection was carried out at a multiplicity of infection (MOI) of 0.3 to ensure that most cells received only one gRNA copy. At 48 hours after infection, cells were transferred to 500 cm^2^ plates and treated with Blasticidin S (4 µg/ml) for 7 days. Five days after infection/transfection, we started treating cells with Zeocin at 100 µg/mL over the course of two weeks to remove cells in which either no donor was inserted, a donor was inserted out of frame, or a product fusion protein was toxic to cells. Surviving cells were then pooled and stored at 500-fold coverage using Cell banker (AMSBIO). One sample of the cells with successful integration was subjected to fluorescent imaging; and for each end (5’- or 3’-) integration, samples were subjected to genomic DNA extraction for posterior analyses.

#### mClover3 Fluorescence Imaging of the pooled library

Forty-eight hours prior to imaging, cells were grown at a density of 7 x 10^5^ cells per dish in 35 mm Nunc™ Glass Bottom Dishes (Thermo Fisher Scientific) in DMEM without phenol red. Cells were imaged on an inverted Nikon Ti2-equipped with 20x/0.5 Air Ph1 WD 2.1, 60x/1.4 Oil DIC WD 0.13, 100x/1.45 Oil Ph3 WD 0.13, 100x/1.45 Oil DIC WD 0.13, 4x/0.2 Air WD 20, 20x/0.75 Air DIC WD 1.0.and 405 nm, 488 nm, 561 nm and 642 nm lasers. Images were taken using 488 nm laser at 50% power, 200ms exposure time, the 60x/1.4 Oil DIC objective, NIS Elements version 4.60 software and analyzed by Fiji software (68).

#### Genomic DNA extraction and genomic PCR

Genomic DNA was extracted from enough cells to maintain coverage per cell line using the Zymo Research Quick-gDNA MidiPrep according to the manufacturer’s protocol. In the next step, all genomic DNA harvested from the screen was PCR amplified using NEBNext high-fidelity polymerase (New England Biolabs) (Supplementary Table 16) and cleaned up using the Zymo-Spin V with Reservoir. NGS was performed on purified samples on Nextseq 150 cycles, Mid Output, v2.5 (Illumina); we aimed for 100 reads per individual gRNA.

#### mClover3 Fluorescence Imaging of single cells

Cells were incubated in a glass bottom 96-well plate (Optical CVG-Thermo Fisher) and were imaged using a GE InCell Analyzer 6000. The imaging parameters were the following: excitation was performed using the 488nm 25mW laser diode at 100% with a 100ms time exposure. Separation between excitation and emission light was performed using a quad band-pass polychromic mirror BP 309/40 + BP 482/18 + BP 564/9 + BP 640/14. Emission light was filtered using a single band pass BP 525/20. One field per well was imaged using a 60x/0.95 Plan Apo air Objective from Nikon. Focus was done using a 785nm laser autofocus with 3µm offset. Images were captured using a 2048x2048 sCMOS camera with 16-bit depth and rolling shutter giving a final pixel size of 0.1083 µm^2^.

#### Image processing of knock-in cells

Images were collected as TIF files. Images were segmented using manual intensity thresholding using ImageJ(68). The mean intensity of the segmented area was measured on the original 16-bit images. Because mClover3 intensity varies greatly between cell lines, brightness and contrast were reset to display the min and max intensities for each image. 8-bit images were then generated and inserted in the figure.

#### Identification of donor DNA insertion site by nested PCR

To identify the location of the donor DNA in the human genome, we used a modified version of a method described by Lourdault, Matsunaga *et al.* to identify the insertion sites of *Leptospira interrogans* transposon mutants in a hamster infection model (38). Here, we adapted their protocol to identify the location of our donor DNA fragments. For this objective, first the cells that have integrated donor DNA fragments were subjected to genomic DNA (gDNA) extraction as described above, and sheared by sonication into fragments of 200-600 bp. Next, we used terminal deoxynucleotidyl transferase (TdT) (New England Biolabs) to add repeats of CCCCCCCCCCCCCCCC (C-tail) at the end of fragmented DNA in accordance with manufacturing protocol. The C-tailed genomic library was subjected to two rounds of PCR. The first PCR was performed with a forward primer which recognized a part of the donor DNA and the reverse primer designed on the C-Tail (GGGGGGGGGGGGGGGG). The product of the first PCR was subjected to the second round of PCR to enable the addition of the required adapters for sequencing. The resulting PCR products were subjected to PCR clean up (Takara) followed by NGS using NextSeq 500 (Illumina) high output 2 x 75 bp Flow Cell.

#### Processing Semi-nested NGS data

Semi-nested PCR NGS data were pre-processed, trimmed, and mapped to the human genome in multiple steps to identify donor DNA integration sites and reading frames. Read processing was performed independently for 5’ and 3’ tagging libraries using Cutadapt v5.0. First, to remove non-informative reads harboring donor plasmid backbone sequences, forward and reverse reads containing full or partial plasmid sequences were filtered out. Next, the tag sequences were trimmed from the 5’ ends of the forward reads using anchored alignment.

To account for potential partial deletions in the donor DNA linker sequence during non-homologous end joining, linker sequences were trimmed by using full length and partial linker as query sequences for Cutadapt. For 5’ libraries, the 36-bp linker sequence was trimmed from the 5’ end of forward reads by incrementally shortening the query sequence down to a minimum of 15 bp, recovering reads with truncated linkers. For C-terminal libraries, a similar incremental approach was applied to the 30-bp linker. Finally, poly-G tails generated during the library preparation were removed from the reverse reads, uncalled nucleotides (Ns) were trimmed from read extremities, and reads shorter than 20 bp were discarded.

The trimmed reads were aligned to the human reference genome (assembly hg38) using Bowtie2 (version 2.3.5.1). Alignments were performed using --local and --very-sensitive-local parameters, with a defined maximum DNA fragment length of 1000 bp (-X 1000). Unaligned reads were discarded, and the resulting SAM files were converted to position-sorted BAM files using SAMtools (version 1.9).

Post-alignment processing and on-target validation were performed using custom Python scripts utilizing the pysam (version 0.23.3) and pandas (version 2.3.3) libraries. To determine if donor DNA integrations were on-target, mapped reads were matched to the nearest expected Cas9 cut site in the sgRNA libraries. Because forward reads sequence directly come out of the donor tag, they precisely define the integration junction and were strictly matched within ± 50 bp of an expected cut site. Reverse reads, which originate from randomly sheared genomic fragments, were permitted a broader 1000 bp tolerance. Reads meeting these distance criteria were considered “on-target”. Finally, for reads containing the junction site of on-target integrations (forward reads), the nucleotide distance between the integration site and the annotated coding sequence (CDS) start or end site was calculated. This distance was used to compute the number of base pairs lost, allowing us to determine whether the donor DNA was integrated in-frame.

## Supporting information

List of gRNAs that were designed to target 18,489 human protein-coding gene sequences at their 5- end.

List of gRNAs that were designed to target 18,489 of human protein-coding gene sequences at their 3- end.

List of gRNAs that were recovered after cloning the gRNA library into LentiCRISPR-V2 to target genes at their 5- end

List of gRNAs that were recovered after cloning the gRNA library into LentiCRISPR-V2 to target genes at their 3- end

List of gRNAs that were recovered after HEK 293T cells infection with the lentiviral library containing guides to target the 5- end of genes

List of gRNAs that were recovered after HEK 293T cells infection with the lentiviral library containing guides to target the 3- end of genes

List of genes that were identified from nested PCR and are tagged with donor DNA at the 5- of human protein-coding gene sequences.

List of genes that were identified from nested PCR and are tagged with donor DNA at the 3- of human protein-coding gene sequences.

List of gRNAs that were recovered after PRISM screening in Hap1, HCT116, and HeLa cells.

List of 38,788 unique accessions for the designed libraries.

List of signal peptide, transit peptide and pro-peptide annotations found at either N- or C- termini for the designed libraries.

List of 34 genes that had no guides in our designed libraries.

## Acknowledgments

The authors thank Professor James Omichinski and Truche Sébastien for providing Cas9 protein used in *In vitro* Cas9 cleavage assay and Monique Vasseur and the Microscopy Platform form the faculty of medicine of Université de Montréal. The authors thank Caitlin Taylor of Illumina Inc. for her support in providing the next-generation sequencing kits used in this study. The authors acknowledge support from Canadian Institutes of Health Research (CIHR) grants MOP-GMX- 152556 (SWM) and RNI00417 (AS) and training fellowships from FRQNT (LG) and Mitacs (MK).

## Author Contributions

Conceptualization: SWM, MK, LG; Methodology-Data Analysis: MK, LG, XCG, MMS, LG, NS, AS. Writing-Review: MK, LG, AS, SWM.

## Data availability

All data generated or analyzed during this study are included in this manuscript (and its supplementary information files).

Scripts for methods referenced in this manuscript are accessible in the GitHub repository: https://github.com/Xavier-Castellanos-Girouard/PRISM_NGS_Analyses.

---

**Supplementary Figure 1.**
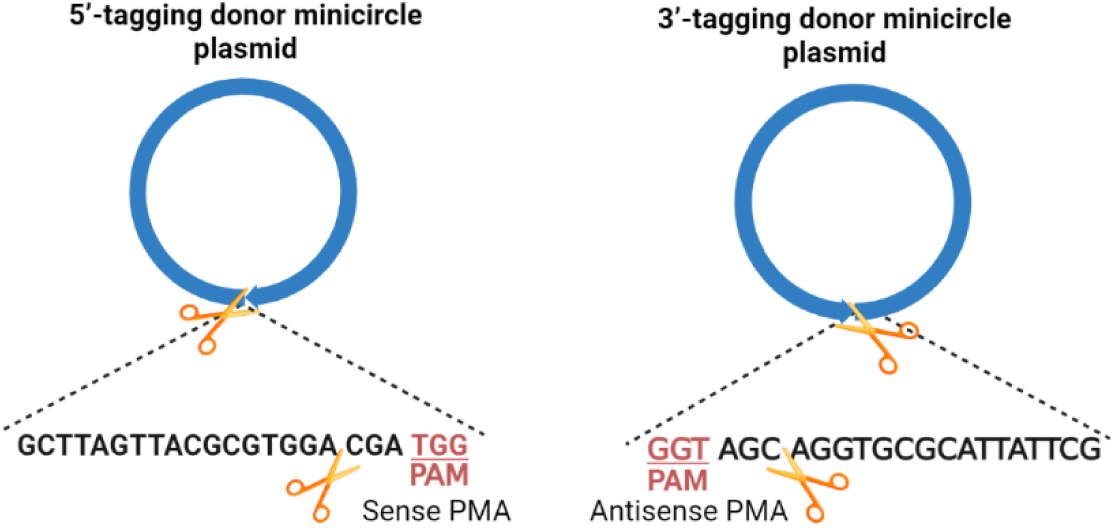
│ MC-Sc-gRNA guide-directed SpCas9-NG PAM sites. The Sc- gRNA was chosen as guide for cleavage of the minicircle (MC) donor plasmids since they do not exist in any protein-coding gene in the human genome. Two types of MC-Sc-gRNA PAM sites were created: donor DNA sequence for 5’ in which the PAM is on the sense strand and donor DNA for 3’ tagging in which the PAM is on the antisense strand.

**Supplementary Figure 2.**
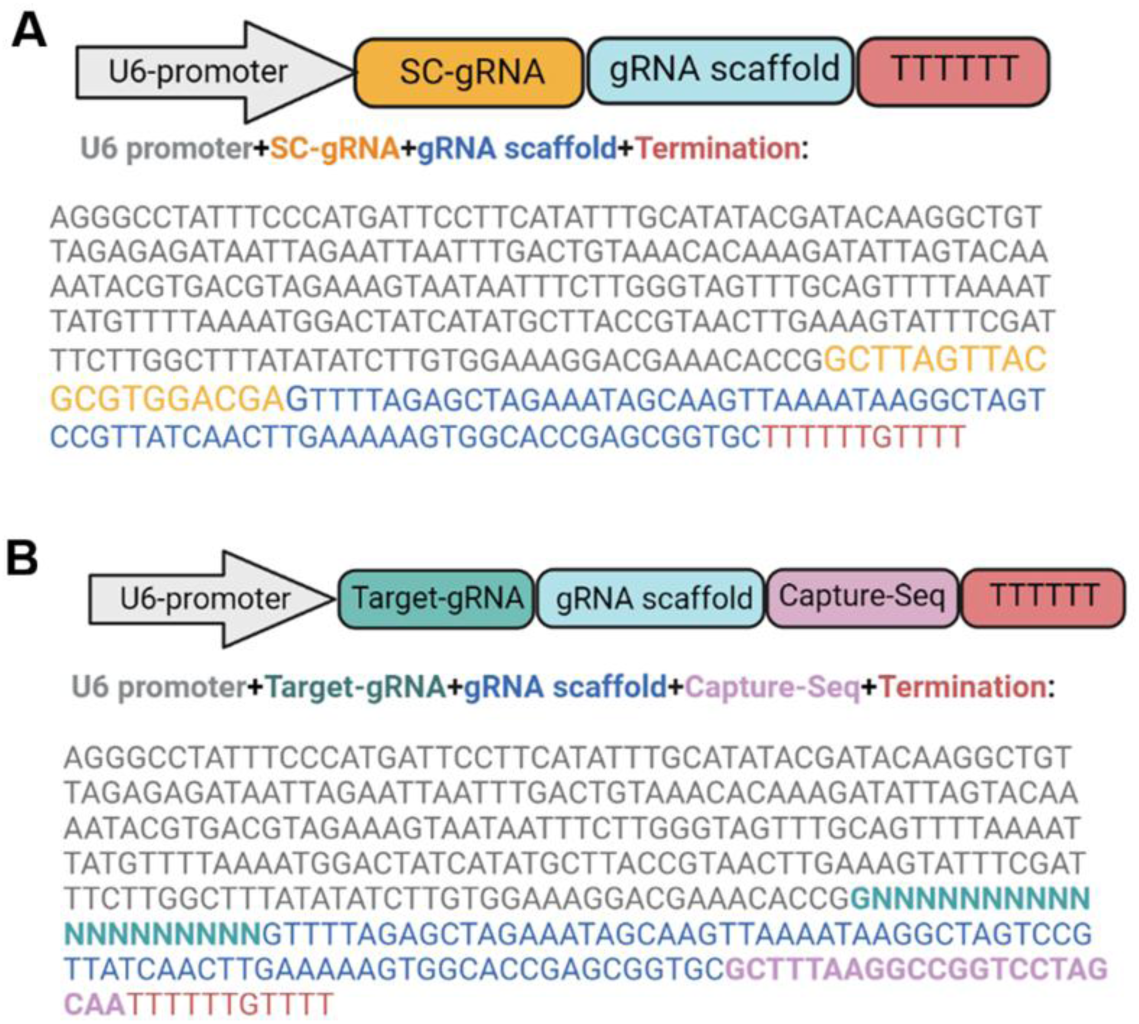
│ Design of gRNA cassettes for donor-cleaving vector and gRNA-CRISPR-Library. a, The scheme indicates the U6 promoter-based expression cassette followed by the Sc-gRNA (orange), gRNA scaffold (blue) and termination signal (red). **b**, The scheme indicates the U6 promoter-based expression cassette followed by the gRNAs targeting 18,489 human genes, including their splice variants (cyan), gRNA scaffold (blue), capture sequence (pink) and termination signal (red). The capture sequence is a 22 bp DNA fragment, designed to provide a priming site to later amplify the gRNA library from the Lenti-CRISPR plasmid and to help to distinguish the donor cleaving-gRNA from the gRNA library in later steps.

**Supplementary Figure 3.**
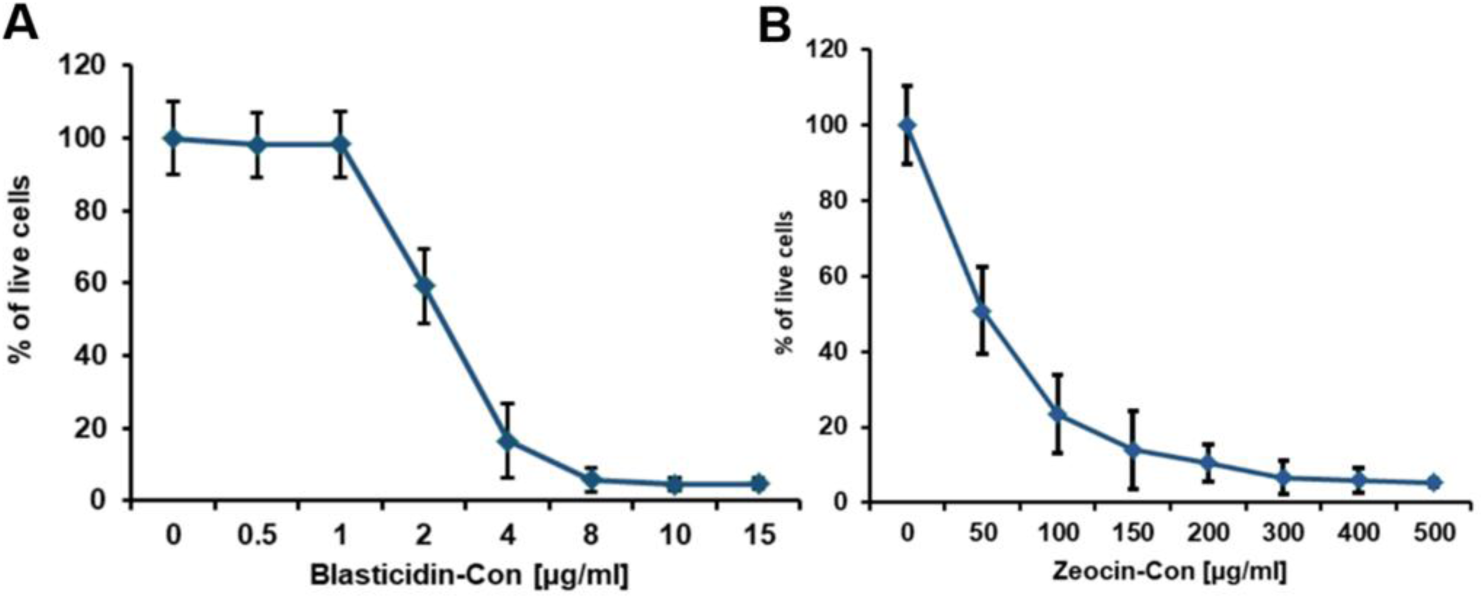
│ Dose response curves for Blasticidin and Zeocin selection for HEK 293T cells. Cell viability assays were performed to determine the minimum amount of the Blasticidin and Zeocin required to eliminate non-infected / non-edited cells over 7 days. HEK 293T cells were seeded in 24-well plates 24 hours prior to antibiotic treatment. In the case of Blasticidin, cells were treated with Blasticidin concentrations between 0 (negative control) and 15 μg/ml; and for Zeocin, cells were treated with concentrations between 0 (negative control) and 500 μg/ml for 7 days. At the end of the screen, cells were treated with Cell Counting Kit-8 (CCK-8, Sigma) according to manufacturer’s instruction; and an estimate of the number of surviving cells was made based on OD_450_ measurements.

**Supplementary Figure 4.**
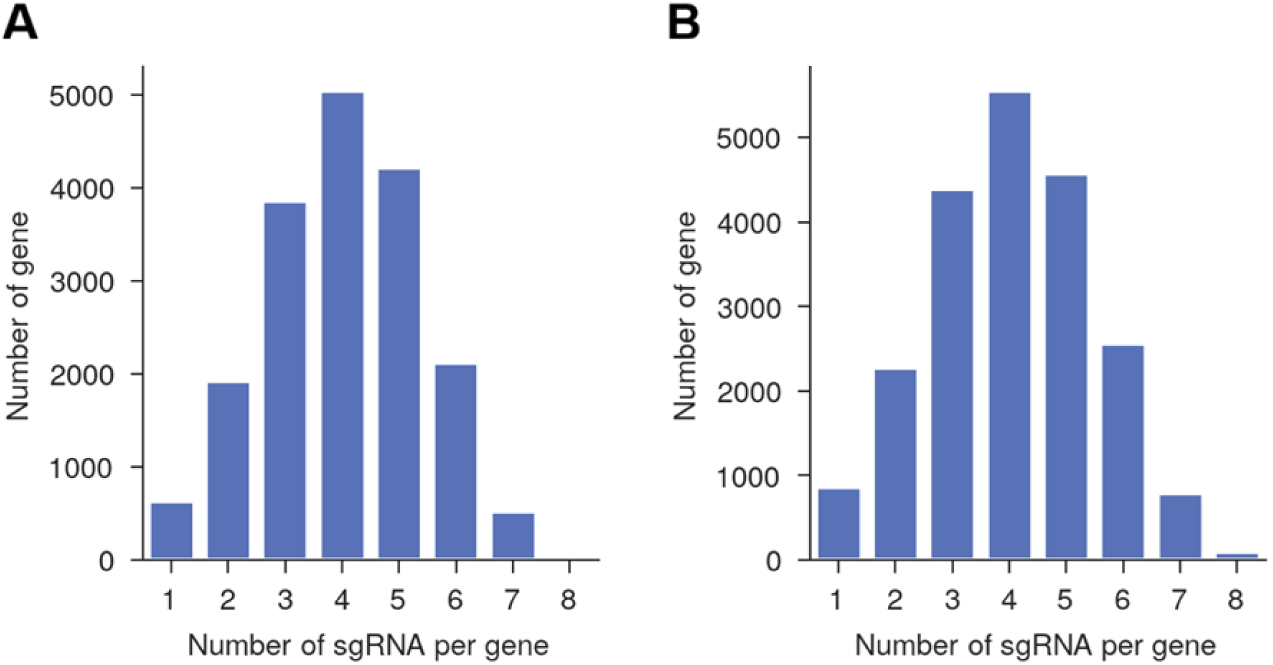
│Design of two genome-wide gRNA libraries consisting of minimum of 1 and maximum of 8 gRNAs per splice variant of human genes. We designed a genome-wide resource of two gRNA libraries consisting of gRNAs that: i, target PAMs resulting only in in-frame double strand breaks; ii, show minimal off-target Cas9 cleavage activity; iii, have maximal on-target Cas9 cleavage activity; iv, do not contain homopolymer stretches (e.g., AAAA, GGGG); and v, meet GC content between 20 to 80%. Following the above criteria, our libraries consist of a minimum of 1 and maximum of 8 gRNAs per splice variant of human genes.

**Supplementary Figure 5.**
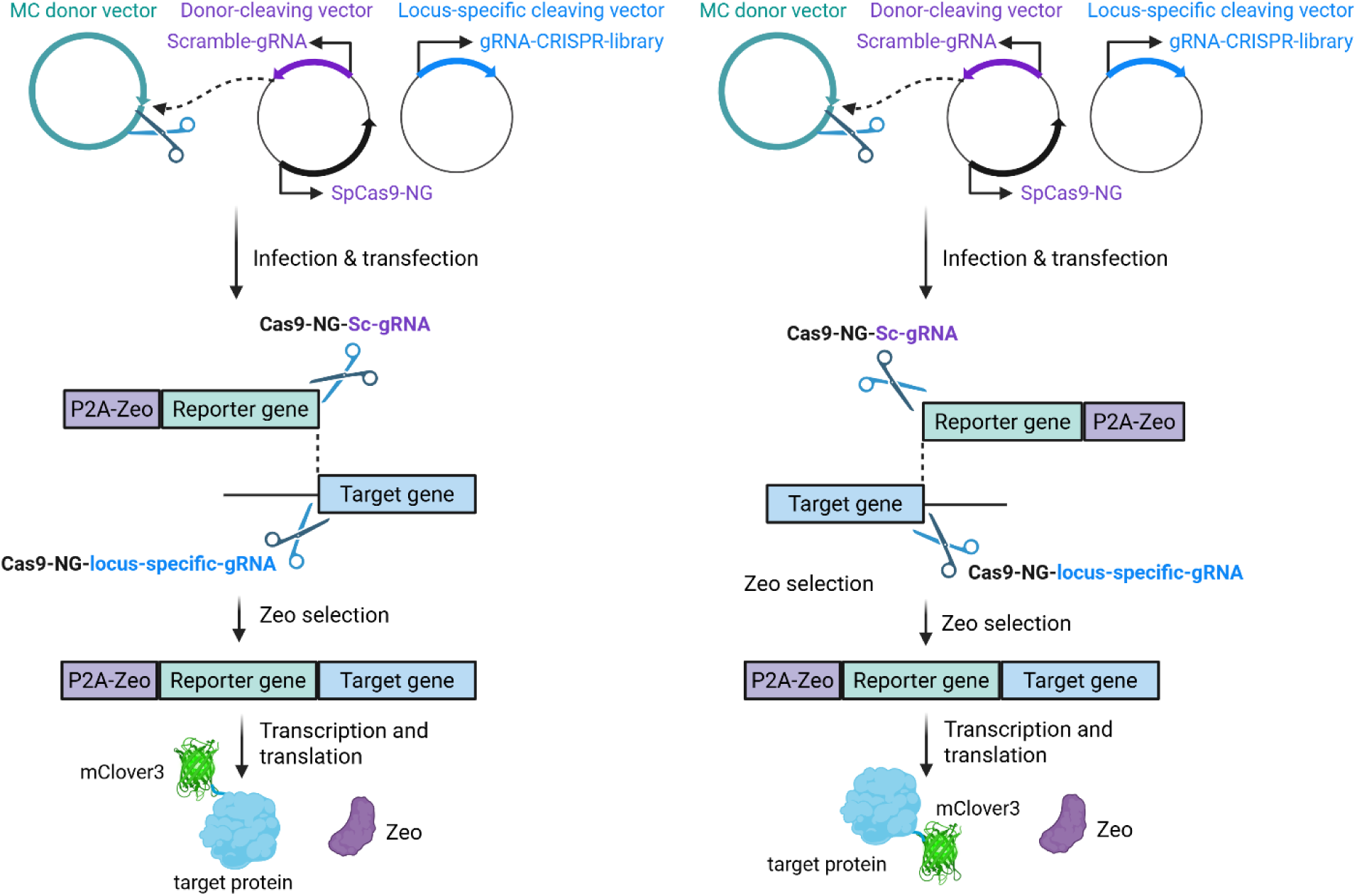
│ Design and assessment of NHEJ-based knock-in modules for integrating mClover3 green fluorescent protein into the 5’ and 3’ end of target protein. Schematic of vectors and methods used to establish knock-in system for gene tagged with mClover3. The system comprises three vectors: a donor vector that contains coding sequences for peptide linker-mClover3 followed by an antibiotic resistance gene as a selection marker fused to P2A self-cleaving peptide (green); a donor cleavage vector that carries Sc-gRNA (purple) and SpCas9-NG (black); and a vector that cleaves a specific target gene in the genome (blue). Created in BioRender.com.

**Supplementary Figure 6.**
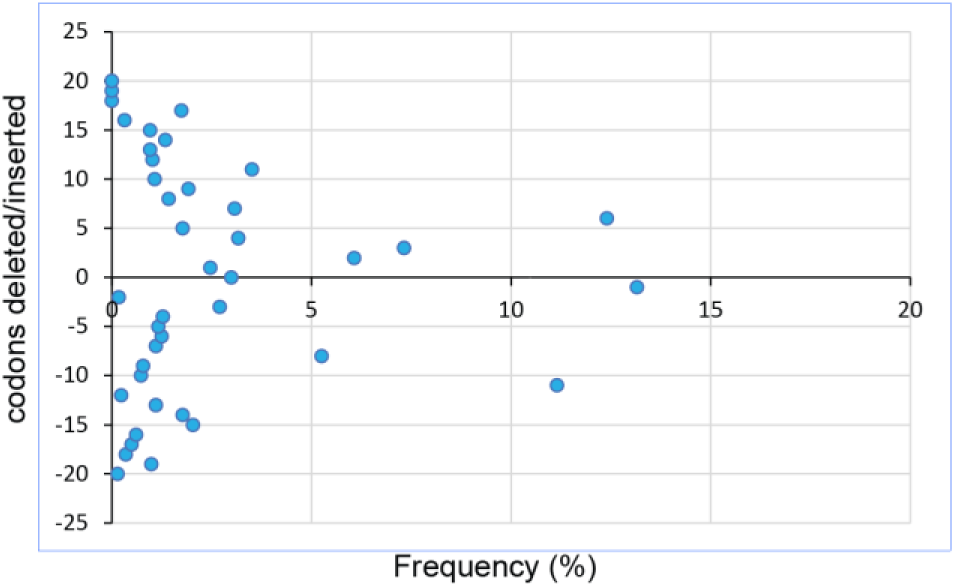
│In-frame indel distribution at integration sites after PRISM knock-in screening. Number of codons deleted/inserted after NHEJ events at targeted integration sites show a broad distribution of frequencies that decreases when moving farther from designed gRNAs cutting sites.

**Supplementary Figure 7.**
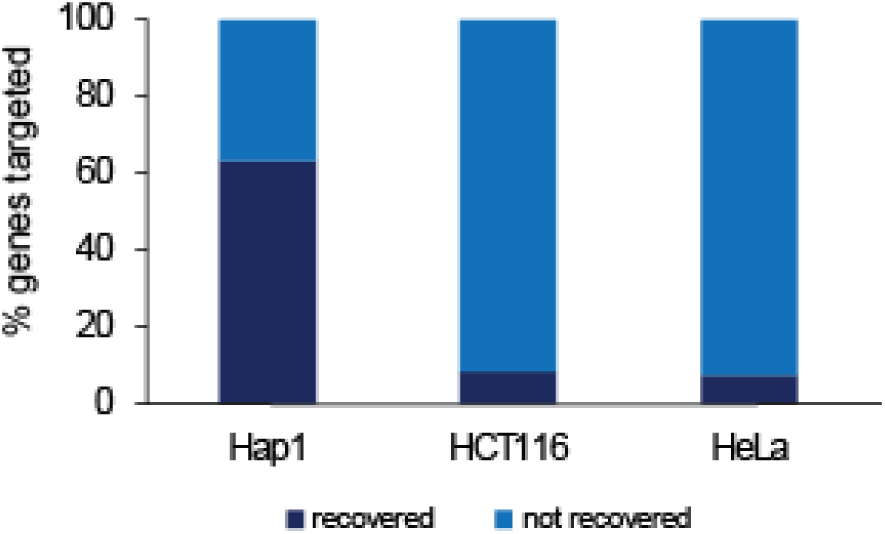
│ Genome-wide CRISPR integration of reporter donor sequence into Hap1, HCT116, and Hela cell lines. Percentage of genes for which guides were retrieved after PRISM knock-in screenings in the cell lines indicated. As shown, the efficiency of the integration changes greatly between different cell lines.

**Supplementary Figure 8.**
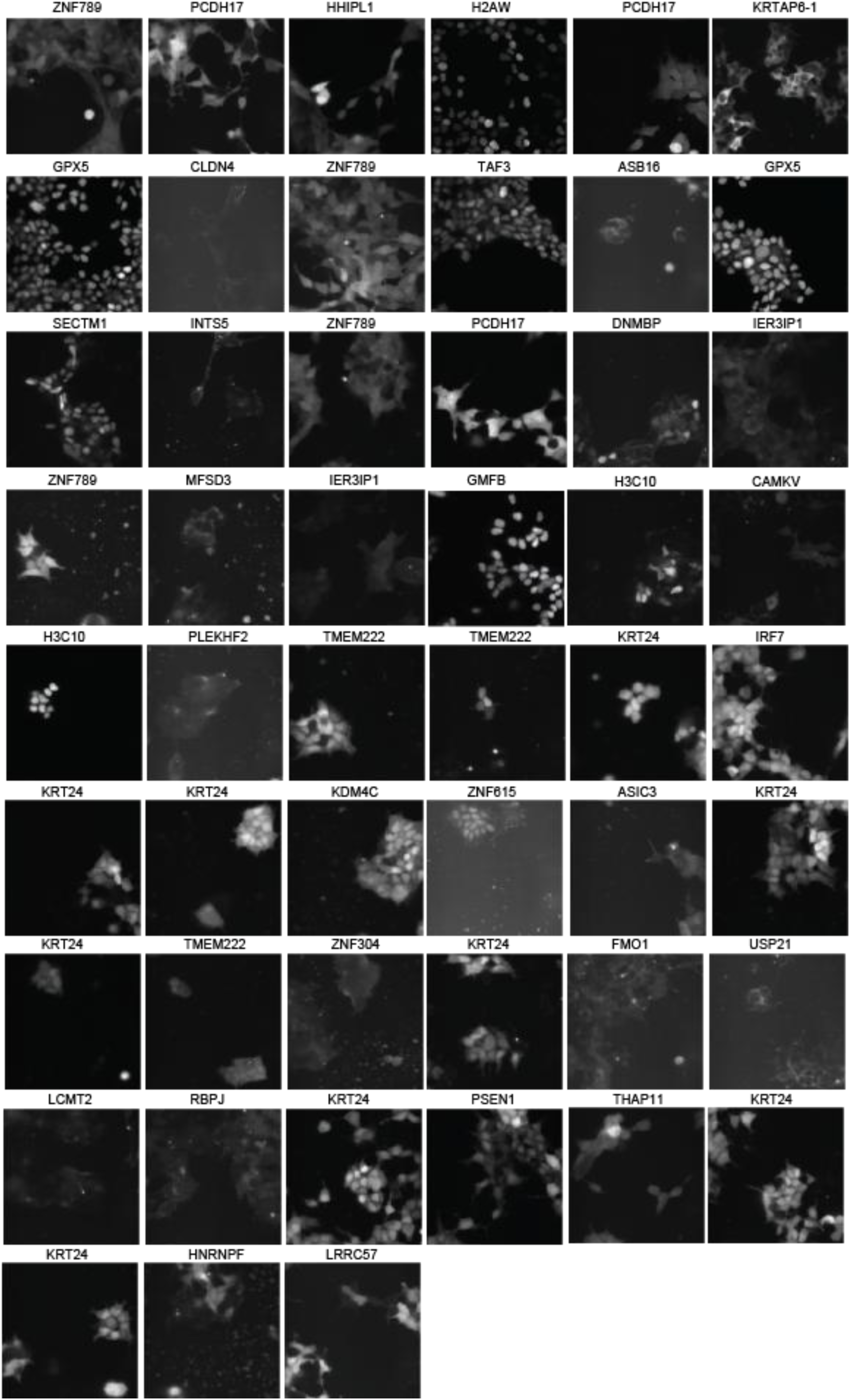
│ Examples of images of successful individual targets showing mClover3 fluorescent signal. Visualization of each single cell in which endogenous proteins are fused to mClover3 using our CRISPR/Cas9-NHEJ-based strategy. We observed variations in fluorescence intensity and patterns of expression, suggesting successful tagging of multiple proteins with varying abundances and different subcellular localizations. Since single cells were collected randomly, some proteins may appear more than once.

**Supplementary Figure 9.**
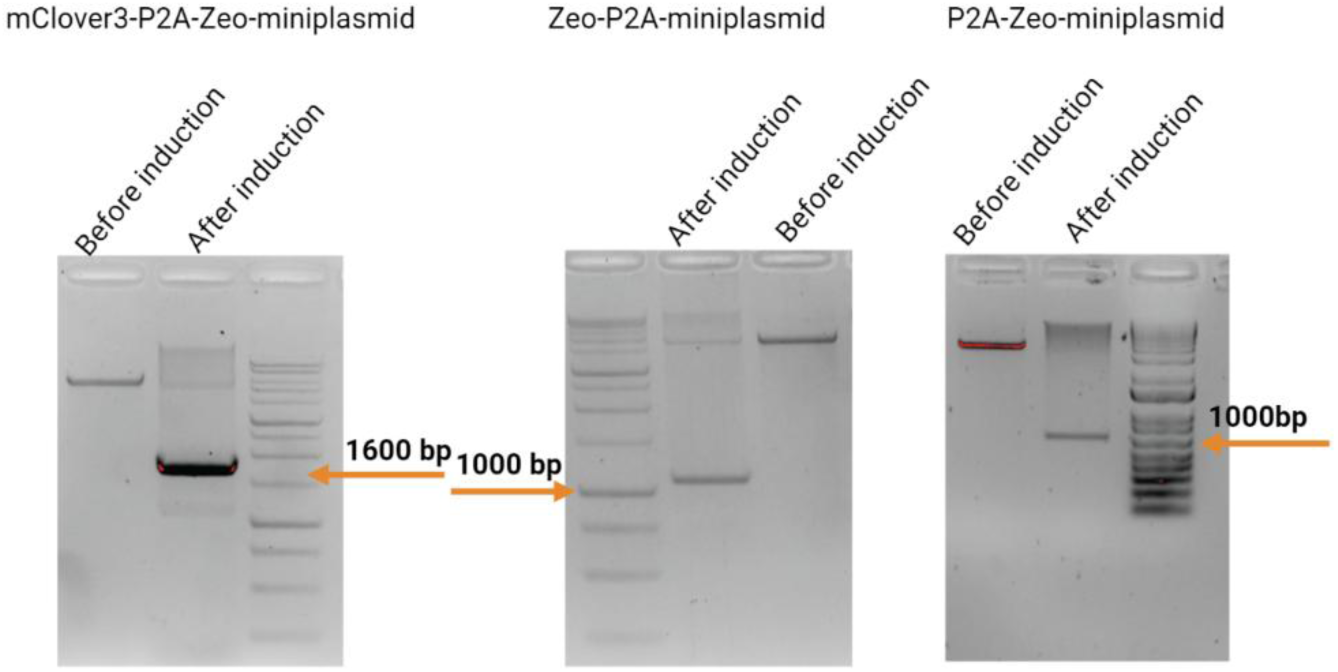
│ Quality of mini plasmid extraction was confirmed by restriction enzyme digestion. 1.0 μg of minicircle plasmid before and after arabinose induction were digested with SmaI restriction enzyme and then run on a 1.5% agarose gel to determine plasmid quality. Parental plasmid indicates the size of plasmid before induction and minicircle shows the plasmid size after induction. Successful minicircle plasmid extraction results in a band at 1600 bp.

**Supplementary Table 1.**
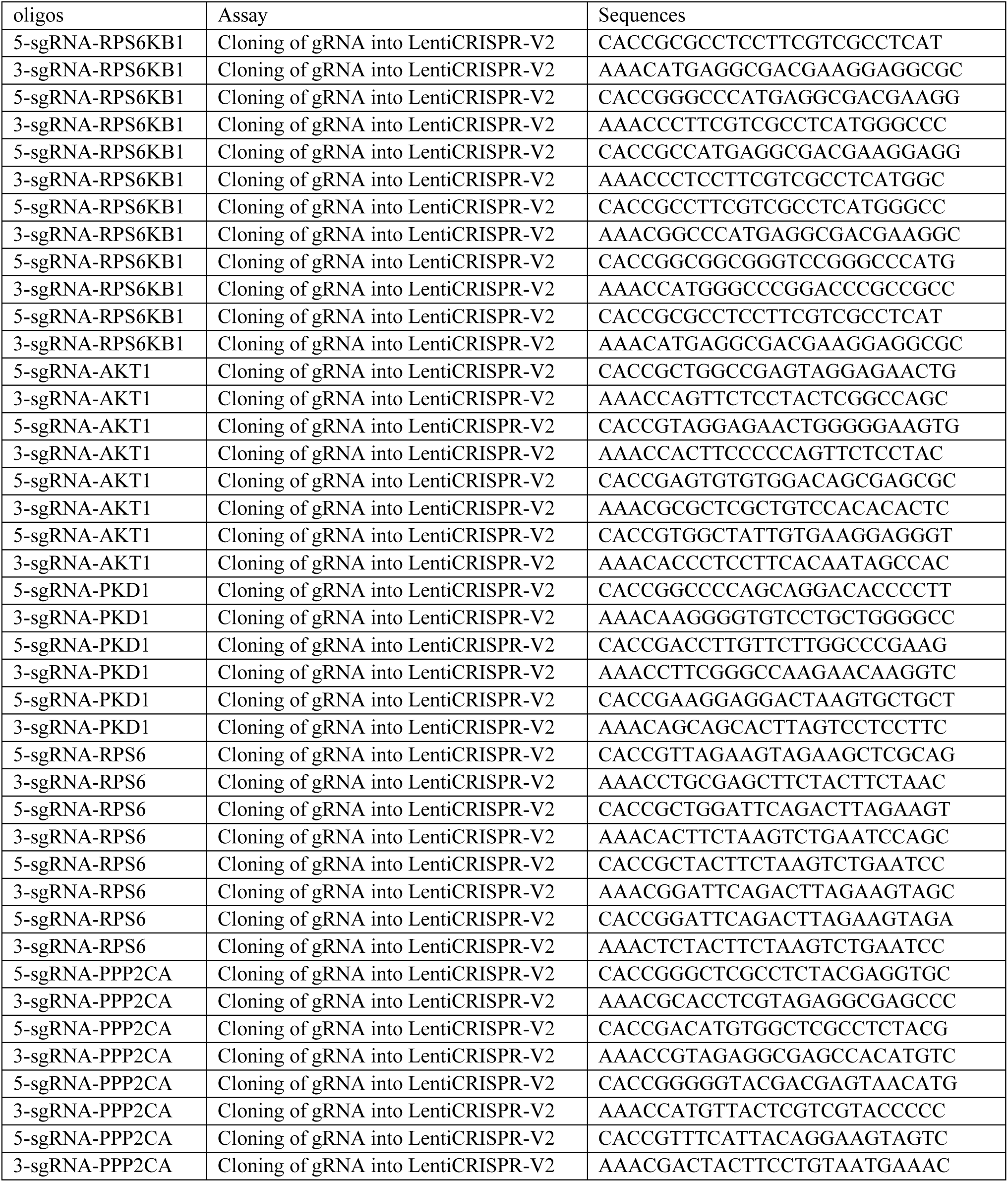
│gRNAs that were designed to target a set of five human-specific genes including RPS6KB1, AKT1, PKD1, RPS6 and PPP2CA at both 5’ and 3’ to protein-coding sequences in HEK 293T cells.

**Supplementary Table 2.** (separate file) │. List of gRNAs that were designed to target 18,489 human protein-coding gene sequences at their 5’ end.

**Supplementary Table 3.** (separate file) │. List of gRNAs that were designed to target 18,489 of human protein-coding gene sequences at their 3’ end.

**Supplementary Table 4.** (separate file) │. List of gRNAs that were recovered after cloning the gRNA library into LentiCRISPR-V2 to target 18,489 of human protein-coding gene sequences at their 5’ end.

**Supplementary Table 5.** (separate file) │. List of gRNAs that were recovered after cloning the gRNA library into LentiCRISPR-V2 target 18,489 of human protein-coding gene sequences at their 3’ end.

**Supplementary Table 6.** (separate file) │. List of gRNAs that were recovered after HEK 293T cells infection with the lentiviral library containing guides to target the 5’ end of human protein- coding gene sequences.

**Supplementary Table 7.** (separate file) │. List of gRNAs that were recovered after HEK 293T cells infection with the lentiviral library containing guides to target the 3’ end of human protein- coding gene sequences.

**Supplementary Table 8.** (separate file) │. List of genes that were identified from nested PCR and are tagged with donor DNA at the 5’ of human protein-coding gene sequences.

**Supplementary Table 9.** (separate file) │. List of genes that were identified from nested PCR and are tagged with donor DNA at the 3’ of human protein-coding gene sequences.

**Supplementary Table 10.** (separate file) │. List of gRNAs that were recovered after PRISM screening in Hap1, HCT116, and HeLa cells.

**Supplementary Table 11.**
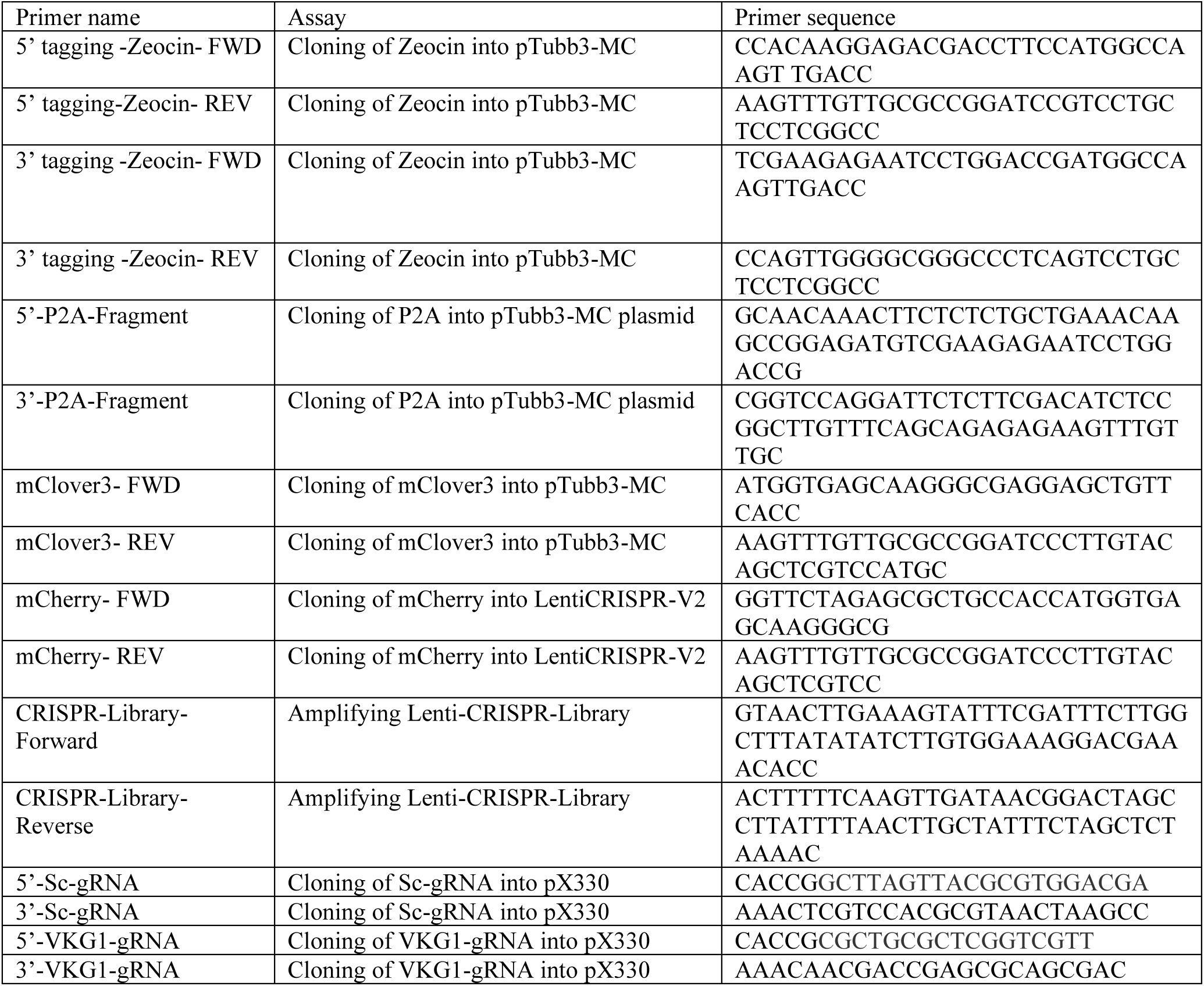
│. Sequences for primers and DNA fragments that were used for preparing MC Donor vectors, donor-cleaving vectors, locus-specific-gRNA-CRISPR-library and preparing DNA templates for *in vitro* transcription of Sc-gRNA and VKG1-gRNA.

**Supplementary Table 12.**
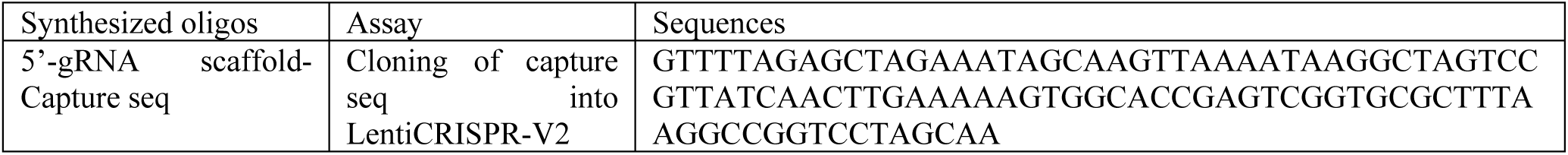
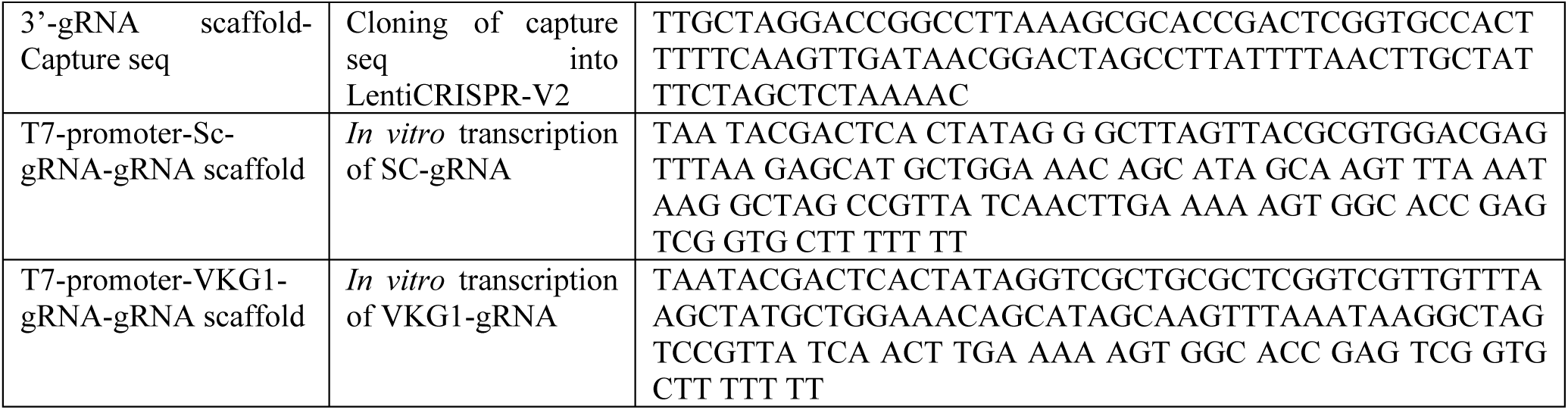
│. Sequences of the synthesized oligos that were used for cloning of Capture Seq after gRNA scaffold and preparing DNA templates for *in vitro* transcription of Sc- gRNA and VKG1-gRNA.

**Supplementary Table 13.** (separate file) │. List of 38,788 unique accessions for the designed libraries.

**Supplementary Table 14.** (separate file) │. List of signal peptide, transit peptide and pro-peptide annotations found at either N- or C- termini for the designed libraries.

**Supplementary Table 15.** (separate file) │. List of 34 genes that had no guides in our designed libraries.

**Supplementary Table 16.**
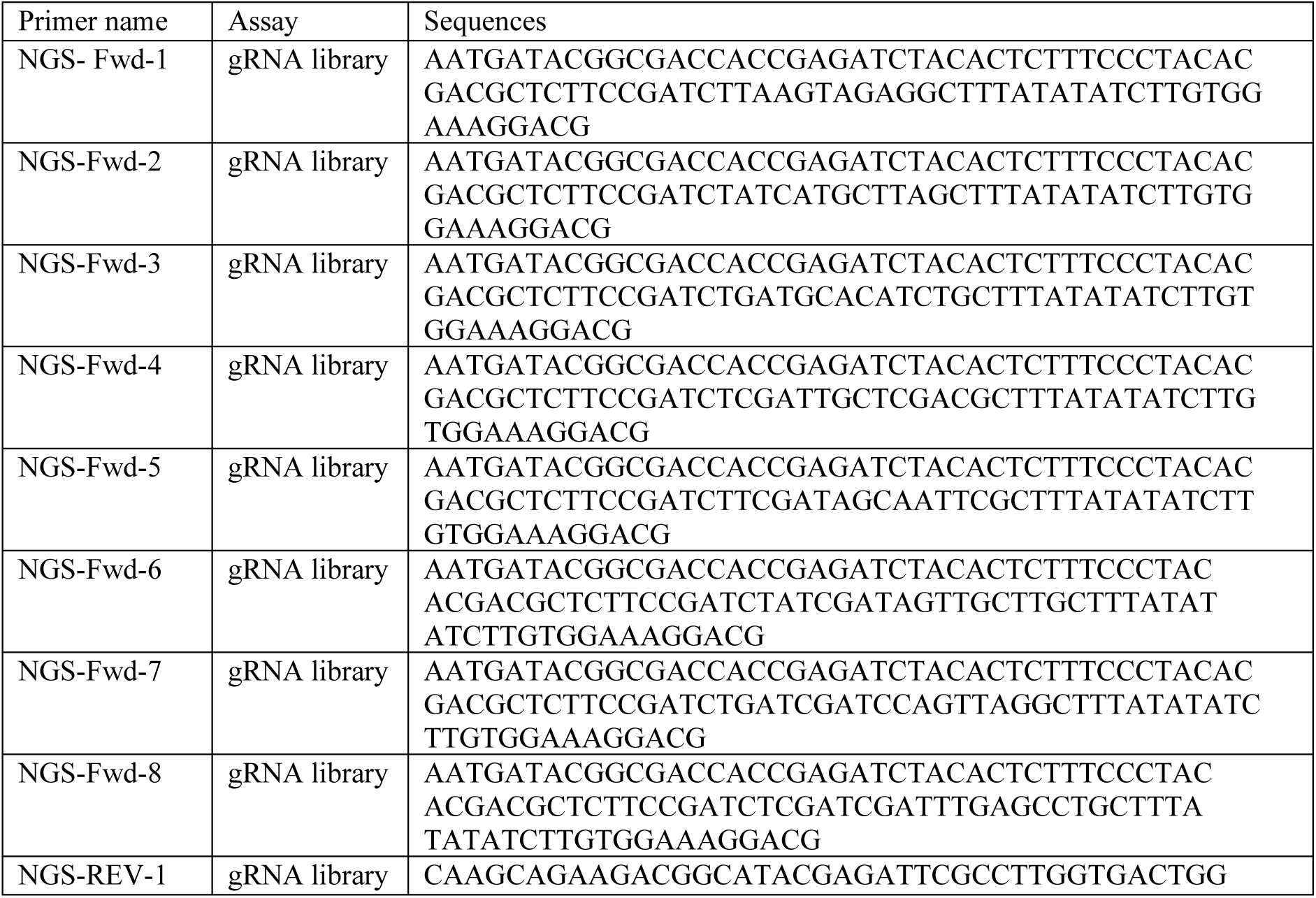
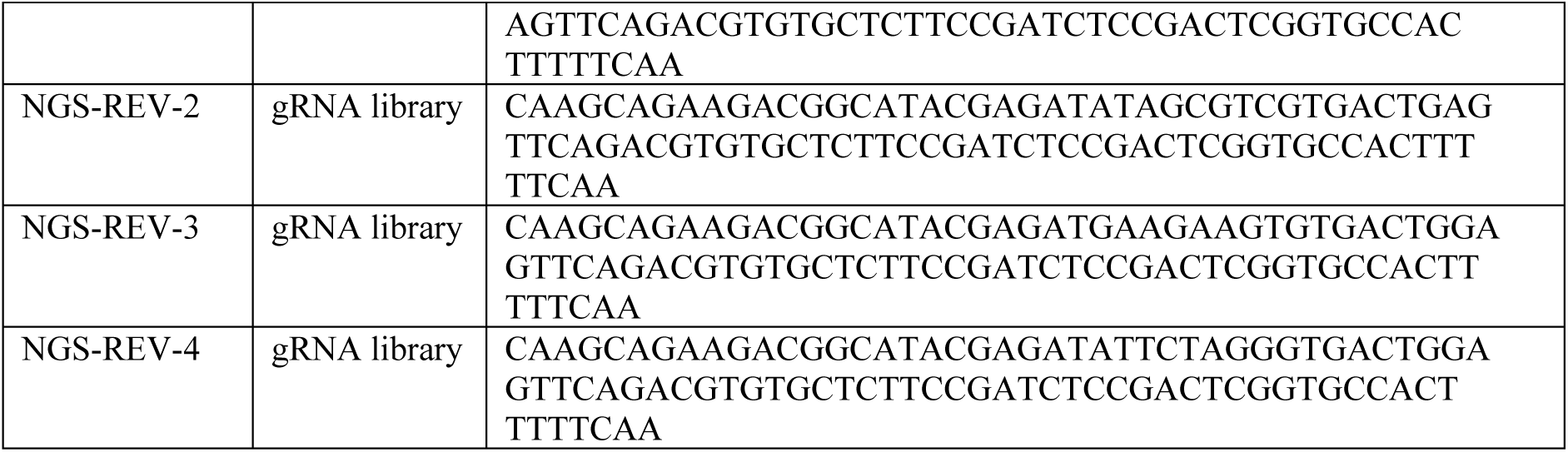
│. Primer sequences for amplifying gRNA-CRISPR-library and NGS.

